# Early intrinsic plasticity of neocortical engram neurons defines memory formation and precision

**DOI:** 10.1101/2024.09.13.612936

**Authors:** Senka Hadzibegovic, Liangying Zhu, Melanie Ginger, Rafael De Sa, Katy le Corf, Maria Gueidão Costa, Yves Le Feuvre, Olivier Nicole, Bruno Bontempi, Andreas Frick

## Abstract

Neocortical memory engrams are thought to stabilize and mature via enhanced interconnectivity during the so-called systems-consolidation process ^1,2^. While synaptic plasticity of these engram connections is considered an important mechanism for storing memories ^3,4^, it cannot fully account for the dynamic vividness of remote, cortically-based memories. Indeed, cell-intrinsic plasticity has been touted as the crucial early priming mechanism that renders nascent engram neurons susceptible to ongoing plastic processes while providing flexibility for later encoding events ^5–7^. Here, we reveal that learning-related neuron-wide intrinsic excitability (IE) plasticity of nascent cortical engram neurons is a permissive mechanism for the formation and specificity of remote associative memories. Using a *c-fos*-dependent genetic and viral system for the targeted labeling of engram neurons in the anterior cingulate cortex (ACC) combined with ex vivo electrophysiology, we found that contextual fear learning triggered a time-dependent increase in their IE signature expressed over days during the early, but not late, phase of memory formation. Remarkably, chemogenetically hyperpolarizing engram neurons during this early plastic phase enhanced their maturation, increasing the strength and context-precision of consolidated memories and preventing memory disturbance caused by an interference event. Altogether, our findings identify *cell-intrinsic* plasticity within nascent ACC engram neurons as an essential tagging mechanism whose features determine the fate and dynamic content of remote memories.

## Main text

Associative memories are gradually embedded within neocortical circuits during systems-level memory consolidation, the process by which remote memories acquire stability and persistence over time ^1,2^. Early (during learning) tagging of the same cortical circuits that will later act as the permanent repository of memory engrams has been identified as a prerequisite for successful consolidation ^8–11^. Cortical memory engrams are thus thought to undergo a time-dependent maturation process from an initially immature (dormant, non-accessible) to an active (retrievable) state ^12,13^. However, the specific processes regulating engram transformation and enduring memory storage remain elusive.

The allocation of neurons to a given memory engram does not occur randomly. Instead, highly excitable neurons within a connected ensemble are preferentially earmarked for integration into an engram ^14,15^. In corollary, learning-induced changes in neuronal intrinsic excitability (IE) could provide a key regulatory determinant for engram formation during memory consolidation. Neuron-wide IE plasticity has been described in several brain regions following learning paradigms ^5–7^. However, while changes in the synaptic strength and connectivity map of engram ensembles are widely viewed mechanisms for storing enduring memories, the role of IE plasticity in this process is unknown. Neuron-wide IE plasticity would tune the responsiveness of that neuron to the majority of its synaptic inputs (i.e., synaptic penetrance ^16^) and prime it to undergo further plasticity (i.e., metaplasticity ^17^), such as in synaptic strength and connectivity. As a result, this mechanism may dictate the fate of engram neurons during systems memory consolidation and permit the establishment of cortical enduring memories. However, neuron-wide IE plasticity would not function as an effective persistent mechanism for storing lasting memories, as it would significantly diminish the storage capacity of these neurons ^5,18^. To fulfill this suggested role, we hypothesized that associative learning tasks activate anterior cingulate cortex (ACC) engram ensembles and trigger neuron-wide IE plasticity in these ensembles as part of the early mechanisms of the memory trace; these changes occur as a consequence of early activation of these cortical neurons, are transient (restricted to the early phase of memory consolidation), and are required as a permissive mechanism to enable the enduring storage of remote memories within the engram network.

### ACC engram neurons are strongly engaged during learning

The prevailing literature suggests a role for the ACC in remote but not early phases of memory formation, making it a central hub within the extended cortical network supporting remote memories ^12,19^. To explore whether these ACC engram ensembles are already tagged by the learning experience (nascent engram neurons) and their functional role in later memory expression, we employed a *c-fos*-dependent labeling approach that combines double-transgenic “Tet-Tag” mice with virus-mediated expression of a fluorescent marker (mCherry) during contextual fear conditioning (CFC) (Fig. 1a). CFC represents a single event that induces an ethologically relevant enduring fear memory, enabling rigorous temporal control of neuronal labeling. In the presence of doxycycline (dox), no labeling was observed following CFC (Fig. 1b). Conversely, CFC performed during a defined dox off time window led to the expression of mCherry in engaged ACC neurons (Fig. 1b). CFC produced robust levels of conditioned freezing in CFC mice tested either 1 day or 30 days following training, compared to context (CTX) only mice, indicating the formation of a durable fear memory (Fig. 1c and 1d; Extended Data Fig. 1).

**Fig. 1.**
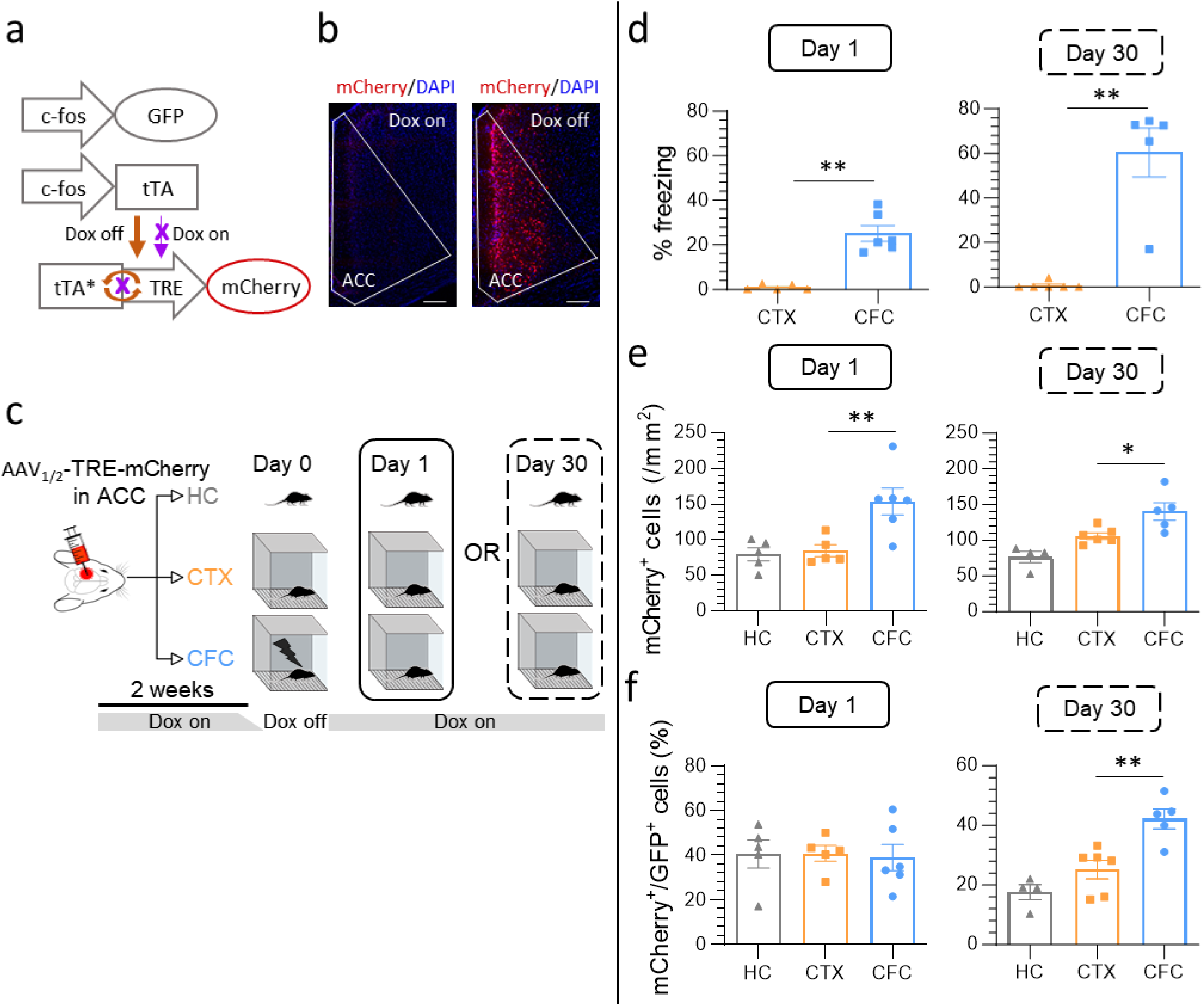
ACC engram ensembles are tagged by learning and reactivated during remote memory expression. **a**, Strategy for long-lasting tagging of cortical engram neurons using the Tet- Tag mouse coupled with AAV_1/2_-TRE-mCherry infusions into the ACC (see panel C). GFP is expressed with a short half-life under the control of the *c-fos* promoter and is used to label neurons active during memory retrieval. Dox removal allows tTA (tetracycline transactivator) expression under the *c-fos* promoter to drive persisting expression of red mCherry protein. **b**, Representative images showing that mCherry-tagging of ACC neurons only occurred in the absence of doxycycline (Dox off). Scale bar: 150 µm. **c**, Experimental design. AAV_1/2_-TRE-mCherry was infused into the ACC of Tet-Tag mice to track the fate of nascent engram neurons. Mice from the CFC (contextual fear conditioning) and CTX (context only) groups underwent training during the dox-off period and were subsequently tested for retrieval either 1 day (recent memory) or 30 days (remote memory) later. Home cage (HC) served as additional baseline controls. **d**, CFC produced robust levels of conditioned freezing in CFC, but not CTX only, in mice tested either 1 day or 30 days following training. **e**, ACC neurons were strongly engaged during CFC as shown by an increase in mCherry^+^ neurons (mice sacrificed one day after CFC) and this tag was stably expressed for at least 30 days. **f**, Reactivation of tagged ACC neurons recruited during initial learning was assessed upon retrieval by plotting the mCherry^+^ to GFP^+^ neurons ratio. Initially tagged (nascent) ACC engram neurons were preferentially reactivated upon remote, but not recent, memory retrieval. Means ± SEMs are shown with individual mouse values. Significance was calculated using Mann-Whitney test (d) or one-way ANOVA (e, f), and indicated in the figures as follows: \**P* < 0.05, \*\**P* < 0.01. See also Extended Data Fig. 1.

Robust CFC-induced fluorescence marker expression was present one day following the learning experience and persisted for at least 30 days, demonstrating rapid and stable labeling of putative ACC engram neurons (Fig. 1e). The number of labeled ACC neurons was significantly greater in the CFC condition compared with those where mice were placed in the conditioning environment without the aversive reinforcement of the foot shocks (CTX) or kept in the home-cage (HC) (Fig. 1e). Early memory retrieval (Day 1) of the CFC group did not significantly reactivate these original putative ACC engram neurons when compared to the CTX and HC groups (similar % colocalization of mCherry^+^ and GFP^+^ neurons; short half-life expression of GFP under the *c-fos* promoter; Fig. 1f). Conversely, these neurons were strongly reactivated during remote memory retrieval (Day 30; increased % colocalization of mCherry^+^ and GFP^+^ neurons; Fig. 1f).

These findings demonstrate that learning tags nascent ACC engram ensembles that are strongly implicated in the expression of remote fear memory. Conversely, these ensembles are not reactivated during the early phases of memory retrieval; rather, they remain in a dormant state.

### Associative learning triggers neuron-wide IE plasticity in ACC engram neurons

Our data show that putative fear-memory-related ACC engram neurons are tagged already during the learning phase. To examine whether this tagging is accompanied by IE plasticity in these neurons, we evaluated their intrinsic properties during the early phase of memory consolidation, one to three days post-CFC. We performed whole-cell recordings of nascent ACC engram neurons (CFC-YFP^+^ neurons, Fig. 2a) within acute ACC brain slices (Fig. 2b), focusing on thick-tufted layer 5 (L5) pyramidal tract (PT) neurons (Fig. 2c), which represent a major type of neocortical output neurons. We probed the plasticity of their intrinsic properties by comparing them with labeled neurons from the CTX group (CTX-YFP^+^ neurons) to identify changes triggered specifically by the learning experience rather than by background activity or unrelated experiences.

**Fig. 2.**
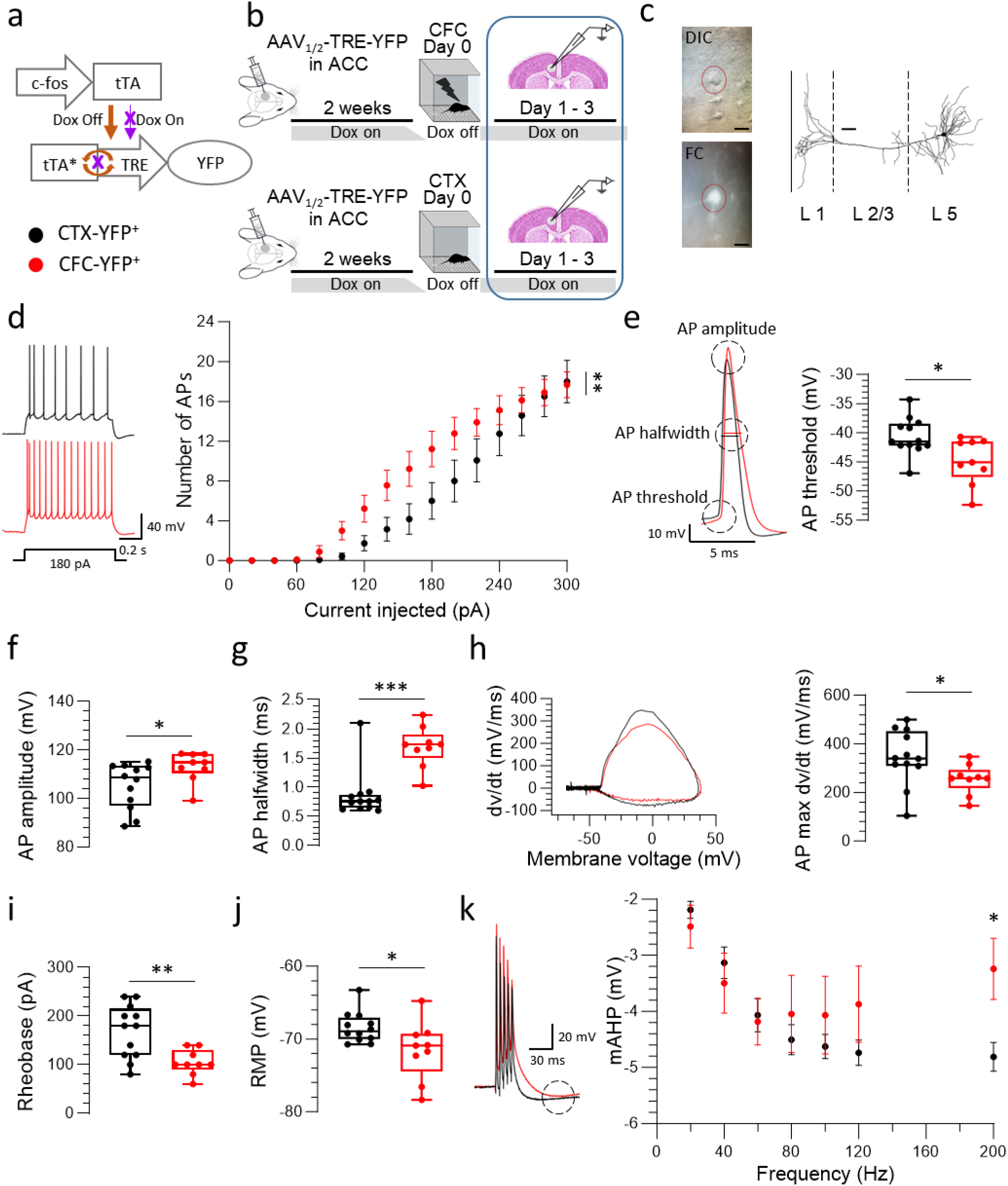
Contextual fear conditioning triggers IE plasticity in ACC nascent engram ensembles. **a**, Strategy for tagging ACC nascent engram neurons for electrophysiological recordings (visualized by YFP expression). **b,** Experimental design. Whole-cell patch-clamp recordings of YFP^+^ neurons were obtained from CFC and CTX groups shortly after training (1-3 days) following training. **c** (left), Representative images from differential interference contrast (DIC) microscopy and fluorescence confocal (FC) image during recording (scale bar 20μm). (right) Three-dimensional reconstruction of a YFP^+^ neuron from layer 5 (L5) of the ACC. Scale bar: 50 μm. **d to m**, Detailed analysis of YFP^+^ neurons from the CFC (n = 9) and CTX groups (n = 12). **d**, (left) Representative traces of action potentials (APs) induced by a 180m pA current injection for a CFC and a CTX neuron. (right) Significant increase in the number of APs as a function of injected current in CFC compared to CTX neurons. **e** (left), Example traces of single APs from a CFC and a CTX neuron, illustrating differences in threshold, amplitude, and half-width. (right) The hyperpolarized AP threshold is higher in CFC neurons. **f, g**, Increase in AP amplitude and AP half-width. **h** (left), Example traces for AP upstroke velocity from a CFC and a CTX neuron (max dv/dt). (right) Max dv/dt is decreased in CFC neurons. **i,** Reduction in rheobase in CFC neurons. **j**, The hyperpolarized resting membrane potential (RMP) is higher in CFC neurons. **k** (left), Example traces of medium after-hyperpolarization (mAHP) from a 5-AP-train at 200 Hz measured from a CFC and a CTX neuron. (right) CFC neurons show a significant reduction in mAHP amplitude for high frequencies. Data points are individual cells with median, lower and upper quartiles, and minimum and maximum data values. Statistical significance was calculated using repeated measures of two-way ANOVA followed by Šídák’s multiple comparisons test (d), mixed-effects model when there were missing values (k) or the unpaired t-test (e-j), \**P* < 0.05, \*\**P* < 0.01, \*\*\**P* < 0.001, \*\*\*\**P* < 0.0001. See also Extended Data Fig. 2.

We discovered that CFC induced a significant increase in the neuron-wide IE of putative ACC engram neurons compared to control neurons (Fig. 2d-k; for a complete list of measures and statistical analysis, see Table 1). This increased excitability was expressed as a greater number of action potentials (APs) triggered by injected current (Fig. 2d). In addition, individual APs were elicited at a lower threshold (Fig. 2e), displayed a higher amplitude (Fig. 2f), were broader (increased half-width; Fig. 2g), and exhibited a reduced upstroke velocity (max dv/dt; Fig. 2h).

In agreement with the hyperpolarized shift in the AP threshold, the minimum current to induce an AP (rheobase) was reduced (Fig. 2i) despite a more hyper-polarized resting membrane potential (RMP) (Fig. 2j). Furthermore, the amplitude of the medium after-hyperpolarization (mAHP) following a train of APs was reduced for the highest AP frequencies tested (Fig. 2k). However, this reduced mAHP amplitude had no obvious impact on the AP adaptation during a train of APs (Extended Data Fig. 2a and 2b). Finally, we also found that the membrane time constant (tau) was prolonged, widening the temporal window for the integration of synaptic inputs (Extended Data Fig. 2c).

Our results show that contextual fear conditioning leads to a plastic increase in the neuron-wide IE of the L5-PT subnetwork of nascent ACC engram neurons, which enhances their responsivity to synaptic inputs. Notably, this IE plasticity is evident 1 day post-CFC and persists for at least three days.

### Neuron-wide IE plasticity is expressed during the early, but not late, phase of memory consolidation

Since neuron-wide IE plasticity is not a suitable permanent storage mechanism, we predicted that this form of plasticity should be transiently expressed during the early, but not late phase, of memory consolidation when memories have matured and are mostly consolidated (>14 days post- CFC). To test this, we assessed the intrinsic properties of CFC-labeled L5-PT neurons at 15-17 days post-CFC (Fig. 3a).

**Fig. 3.**
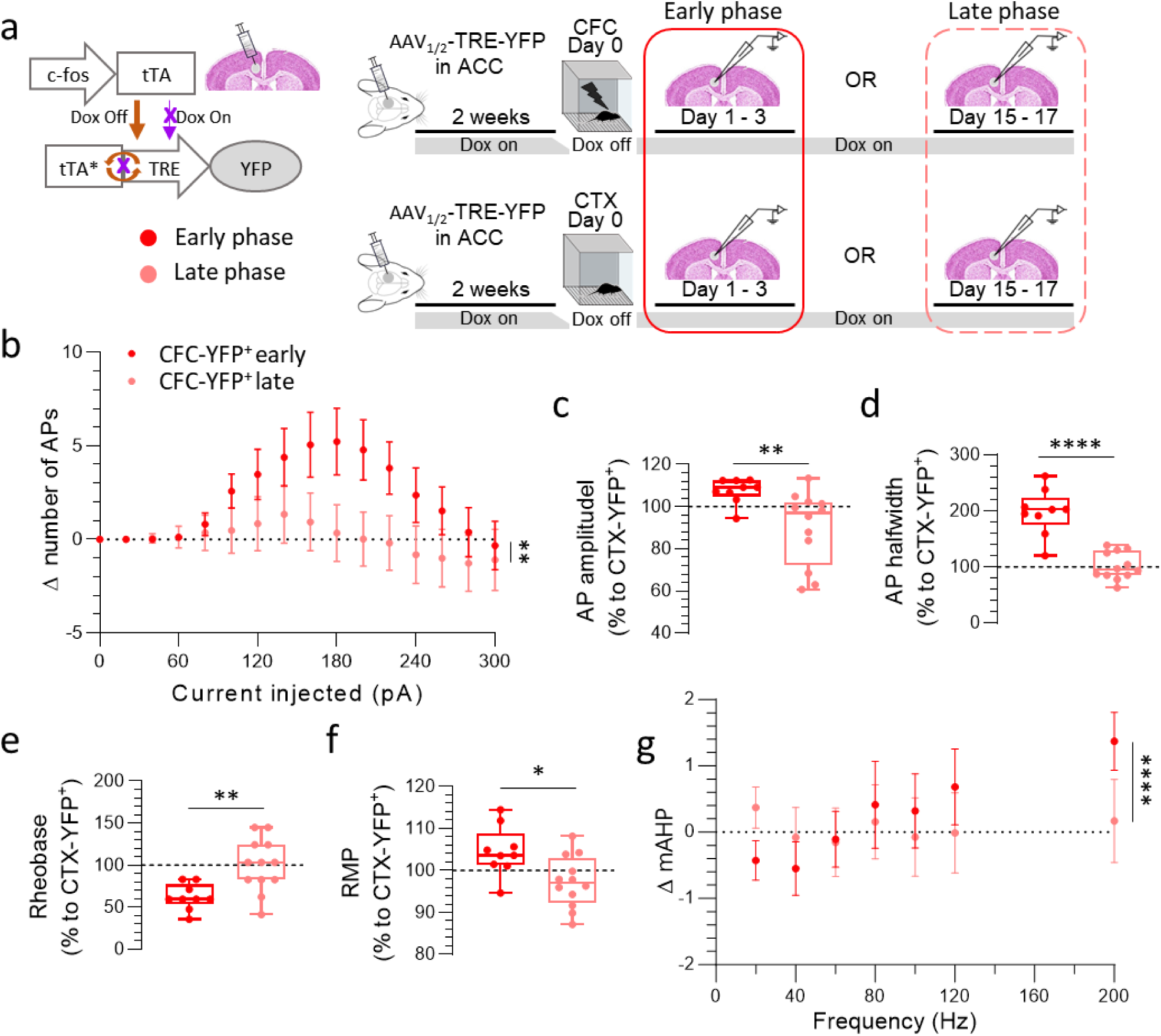
Learning-induced IE plasticity of ACC engram neurons is transient. **A** (left), Strategy for tagging of ACC engram neurons visualized by YFP neuronal expression. (right) Experimental design. Whole-cell patch-clamp recordings were performed from tagged (YFP^+^) L5 pyramidal neurons either during the early phase (1-3 days post-CFC) or late phase (15-17 days post-CFC) of memory consolidation. Data are normalized to values from control (CTX-YFP^+^) cells for the respective consolidation phases. **b-g**, Detailed analysis of YFP^+^ neurons from the CFC groups for each consolidation phase (early phase, n = 9 cells; late phase, n = 12 cells). Significant differences were found for the number of APs (presented as Δ: difference in the values between CFC-YFP^+^ and CTX-YFP^+^) (**b**), AP amplitude (**c**), AP halfwidth (**d**), rheobase (**e**), RMP (**f**) and mAHP amplitude, presented as Δ: difference in the values between CFC-YFP^+^ and CTX-YFP^+^ (**g**). Data points are individual cells with median, lower and upper quartiles, and minimum and maximum data values. Statistical significance was calculated using repeated measures of two-way ANOVA followed by Šídák’s multiple comparisons tests (b), mixed-effects model when there were missing values (g) or the unpaired t-test (c-f), \**P* < 0.05, \*\**P* < 0.01, \*\*\**P* < 0.001, \*\*\*\**P* < 0.0001. See also Extended Data Fig. 4.

These measures confirmed our prediction, showing that neuron-wide IE plasticity has returned to control values (neurons tagged in the home cage) during the late memory consolidation phase. Thus, at 15–17 days post-CFC, the IE features were similar between putative engram ensembles (CFC-YFP^+^) and control neurons (CTX-YFP^+^) of the ACC (Extended Data Fig. 3a-l, Table 2). Accordingly, the following IE measures of putative engram ensembles (CFC-YFP^+^) differed significantly between the early and late phases of memory consolidation (values were normalized to those of CTX during early and late phases, respectively; Fig. 3b-g and Extended Data Fig. 4a-f; for a complete list of measures and statistical analysis see Table 3): AP number triggered by current injection (greater during the early consolidation phase; Fig. 3b), AP peak amplitude (increased; Fig. 3c), AP half-width (increased; Fig. 3d), rheobase (reduced; Fig. 3e), RMP (more depolarized; Fig. 3f), mAHP amplitude (reduced; Fig. 3g), AP adaptation (reduced; Extended Data Fig. 4b), and tau (increased; Extended Data Fig. 4c). Conversely, there was no significant difference in the input resistance (Extended Data Fig. 4d), AP upstroke velocity (max dV/dt; Extended Data Fig. 4e), and only a trend for the AP threshold (more hyperpolarized during the early phase; Extended Data Fig. 4f).

Collectively, these data show that the learning-induced neuron-wide IE plasticity of putative ACC engram neurons is expressed during the early, but not late, phase of memory consolidation.

### The IE state of ACC nascent engram neurons is crucial for the formation of remote fear memory

According to our hypothesis, neuron-wide IE plasticity is expected to enhance the reactivation of the engram ensemble and permit more enduring plastic changes that require repeated neuronal activation and more time to develop. Consequently, modulating the excitation state of ACC engram ensembles during the CFC-induced IE plasticity phase should interfere with this functional role and impact the formation of enduring fear memories. We probed this by using a pharmacogenetic approach. Employing our tet-tagging systems, we expressed either inhibitory (i) or excitatory (e) designer receptors exclusively activated by designer drugs (DREADD) selectively in nascent ACC engram neurons following CFC (Fig. 4a). Mice were then treated twice a day for seven consecutive days following CFC with the DREADD ligand, CNO, or vehicle (saline), and finally tested for remote fear memory expression (30 days post-CFC; Fig. 4a). Surprisingly, we found that chemogenetic hyperpolarization of ACC engram ensembles (including the L5-PT subnetwork) during the early consolidation phase enhanced remote fear memory expression (manifested as increased freezing) in these mice (iDREADD-CNO group) when compared to control (i/eDREADD-saline group; Fig. 4b). In contrast, chemogenetic depolarization of tagged ACC engram ensembles reduced remote fear expression (eDREADD-CNO group; treatment x context interaction effect; Fig. 4b).

**Fig. 4.**
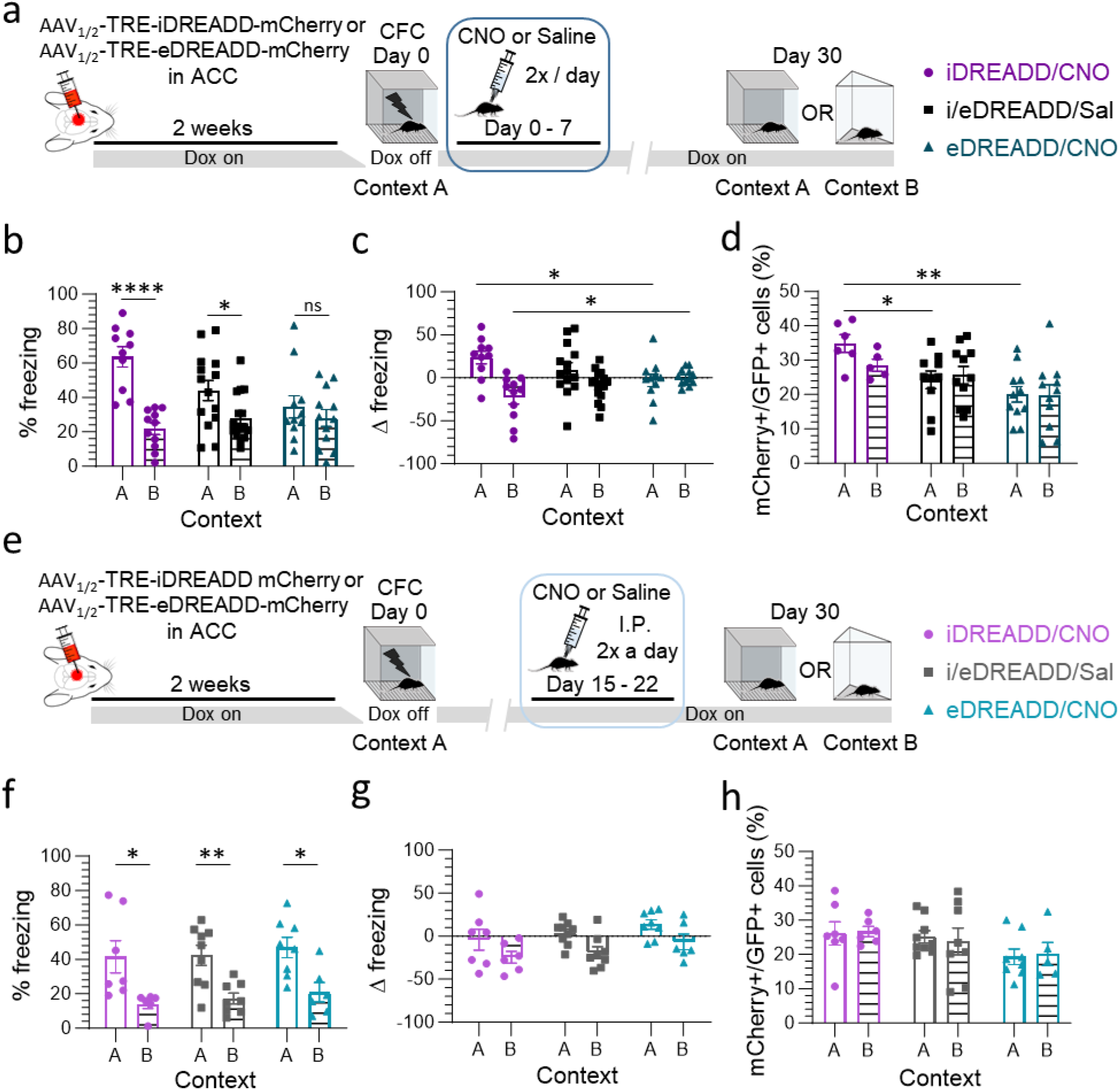
Modulating IE of nascent ACC engram neurons determines the formation and precision of remote fear memories. **a**, Experimental design. Assessing the consequence of manipulating the IE of ACC engram ensembles during the early phase of memory consolidation on context precision of remote fear memories (day 30 post-CFC). **b,** Results for the three groups (iDREADD/CNO, eDREADD/CNO, and i/eDREADD/Sal) are expressed as the % of time spent freezing during remote memory retrieval. **c,** Results are presented as Δ freezing (difference in the % of freezing between retrieval and after the last shock during memory acquisition). **d,** % of reactivated tagged ACC engram neurons following remote memory retrieval (mCherry^+^/GFP^+^ cells). **e,** Experimental design. Same as in (**a**) with IE manipulation delivered during the late phase of memory consolidation. **f-g**, Targeting the late phase of memory consolidation did not alter remote fear memory precision (all groups were able to discriminate context A from context B) as measured by % freezing (**f**) and no changes were observed for Δ freezing (**g**), in either iDREADD/CNO or eDREADD/CNO mice compared to the i/eDREADD/Sal group. **h**, % of reactivated tagged ACC engram neurons during remote memory retrieval. Means ± SEMs are shown with individual mouse values. Statistical significance was calculated using two-way ANOVA followed by Tukey/Šídák’s multiple comparisons test (b-d, f-h), \**P* < 0.05, \*\**P* < 0.01, \*\*\*\**P* < 0.0001. See also Extended Data Fig. 6 and 7.

These data demonstrate that pharmacogenetic hyperpolarization of ACC engram neurons during the early phase of memory consolidation when they display increased neuron-wide IE can enhance enduring fear memory formation.

### Context specificity of remote fear memory is malleable during the early IE plasticity phase

In addition to strengthening remote fear memory, we wondered whether modulating the excitability of ACC engram ensembles during the early phase of memory consolidation could also improve the precision of the remote fear memory for the specific context. We assessed precision by comparing memory specificity for the original context (Context A) versus a new context (Context B) (Fig. 4a). Instead of repeated testing, mice were divided into separate groups for remote retrieval in either Context A or Context B because we found that successive retrieval in both contexts attenuated fear expression in the second context (Extended Data Fig. 5a-c). These experiments showed that the iDREADD/CNO group discriminated better between the two contexts compared to the control group, displaying an improved Context A-specific fear memory (Fig. 4b and c). The improvement in memory precision resulted from an increase in freezing behavior in Context A, coupled with a decrease in freezing behavior in Context B (Fig. 4c). In contrast, activating eDREADD in tagged ACC engram neurons eliminated the capacity to discriminate between the original and the new context (Fig. 4b and c). The differences in memory precision and strength between the three groups could neither be explained by the original freezing score during CFC (Extended Data Fig. 6a and b), nor by the number of engram neurons that expressed DREADD (Extended Data Fig. 6c).

We then asked whether the increased memory strength/precision induced by iDREADD-mediated hyperpolarization of ACC engram neurons (Fig. 4b and c) would be reflected in a greater reactivation of these neurons during remote memory retrieval. Indeed, the reactivation ratio (% colocalization mCherry^+^/GFP^+^ neurons) was significantly greater compared to the saline or eDREADD condition (Fig. 4d). In contrast, the number of retrieval-activated neurons itself was not altered by chemogenetic manipulation (Extended Data Fig. 6d).

Our results suggest that the IE plasticity of nascent ACC engram ensembles during early memory consolidation is crucial for the permanent storage and precision of fear memories. Any modulation of the excitation state of ACC engram ensembles during the IE plasticity phase crucially influences these memory features in a bidirectional manner; chemogenetic hyperpolarization effectively improves memory strength and specificity and results in an enhanced maturation of the nascent ACC engram ensemble, whereas chemogenetic depolarization has the opposite effect.

### Modulating the IE state of matured ACC engram neurons does not affect memory expression

We next examined whether the effectiveness of manipulating the excitation of ACC engram neurons to improve memory formation was dependent on the phase of memory consolidation. To this end, we delayed the DREADD-induced excitability modulation of ACC engram neurons by administering CNO 15 to 22 days following CFC. This timing allowed us to selectively target the late phase of memory consolidation when the IE plasticity had returned to control levels (Fig. 3). Consistent with the previously observed temporal course of IE plasticity, we observed no functional consequence for memory performance or context precision upon remote memory retrieval by either iDREADD or eDREADD activation during this remote time window (Fig. 4f and g and Extended Data Fig. 7a-d). Likewise, ACC engram ensemble reactivation was not affected by excitability modulation at this late phase (Fig. 4h).

Our data shows that the time course of the IE plasticity of ACC engram ensembles coincides with the efficiency of excitability manipulation for fear memory formation and precision, revealing that the remote phase of the consolidation process has become resistant to such manipulation.

### The impact of interference on memory performance depends on the consolidation phase

Interference occurring shortly after training, during the early phase of memory consolidation, has been shown to impair remote memory formation, likely by reallocating more excitable engram neurons into a new ensemble ^14^. Concurring with this, we found that retroactive interference in the form of exposure to an enriched environment 1 day following CFC (Early–INT; Fig. 5a) impaired memory formation by decreasing freezing during remote memory retrieval compared to the control condition (day 30; Fig. 5b). In contrast, the same interference experience administered during the late phase of memory formation (15 days post-CFC; Late–INT) had no impact on remote memory performance (Fig. 5a and b). Freezing levels during CFC were similar between the different groups, ruling out differences in memory acquisition as a potential confounding factor (Extended Data Fig. 8a and b).

**Fig. 5.**
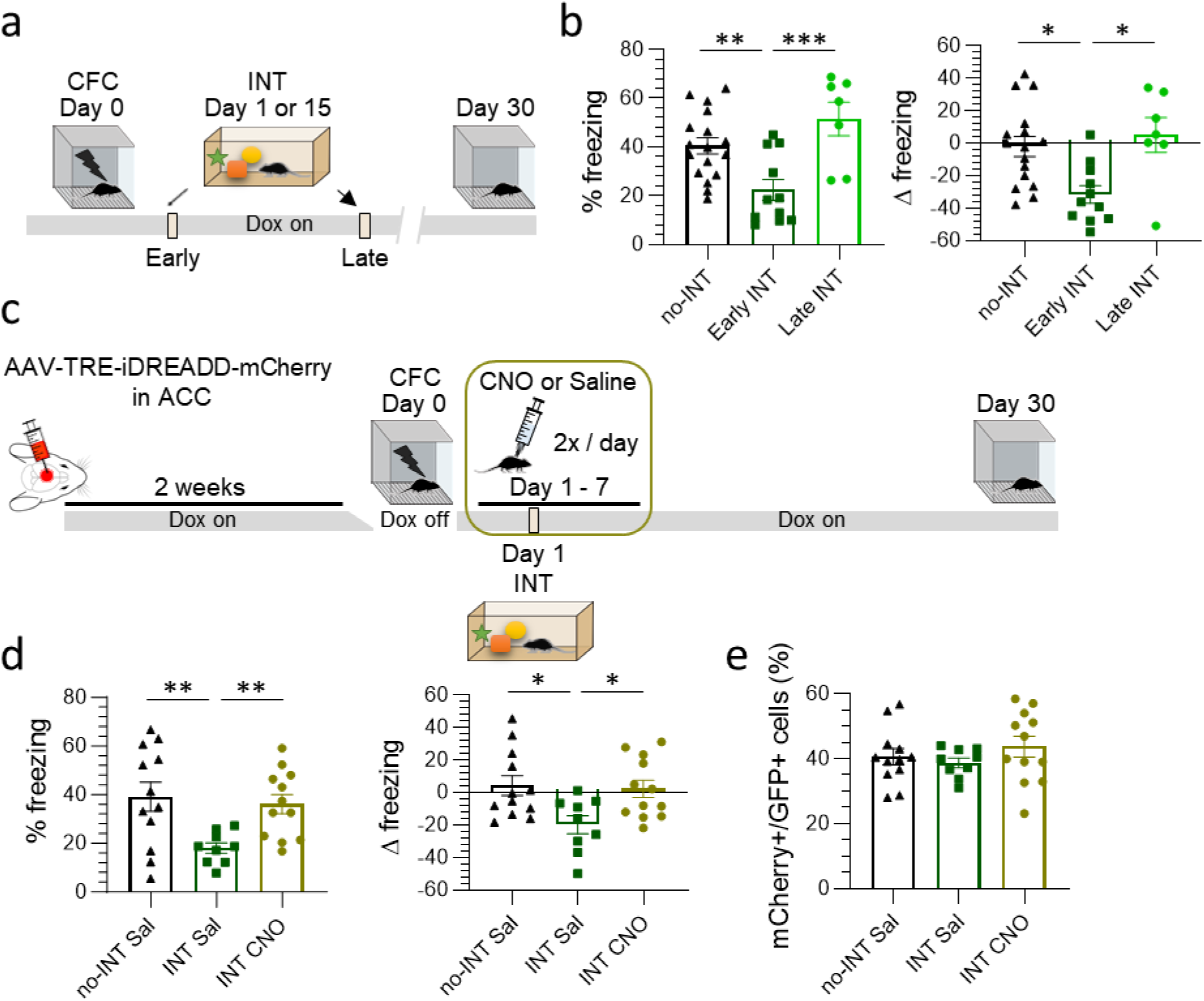
Early manipulation of the IE state of ACC engram neurons prevents memory impairment caused by interference. **a**, Experimental design for evaluating remote memory expression following interference (INT, exposure to an enriched environment). Mice underwent interference either one day or 15 days post-CFC. **b**, Difference in fear expression (% freezing and Δ freezing) between the three groups (no interference, early interference, late interference). **c,** Experimental design. Mice from the iDREADD group were injected with CNO for 7 days post- CFC, with INT occurring 24 h following CFC and memory retrieval being tested on day 30. **d**, Differences in fear expression between the three groups are shown. **e,** percentage of reactivated (ratio mCherry^+^/GFP^+^ cells) ACC engram neurons during remote memory retrieval. Means ± SEMs are shown with individual mouse values. Statistical significance was calculated using the unpaired t-test (b, d, e), \**P* < 0.05, \*\**P* < 0.01, \*\*\**P* < 0.001. See also Extended Data Fig. 8.

These data demonstrate that, similarly to the DREADD manipulation, the impact of an ethologically relevant experiential factor such as interference on memory formation phase-locks to the transient time course of neuron-wide IE plasticity of ACC engram ensembles.

### Precocious manipulation of IE plasticity protects memory formation from interference

We next asked whether chemogenetic hyperpolarization of ACC engram neurons during the early phase of memory consolidation could prevent remote memory impairment caused by an early interference experience. To address this, we administered CNO for the first seven days following CFC to activate iDREADD on ACC engram neurons, exposed the mice to interference on day one, and evaluated remote (30 days post-CFC) memory performance (Fig. 5c). Importantly, this excitability manipulation prevented the interference-induced memory impairment, rescuing the fear expression levels upon retrieval (Fig. 5d). Freezing levels during CFC were similar between the different groups indicating that there were no differences in memory acquisition that might impact these results (Extended Data Fig. 8c and d). The size of the putative engram that supported the acquisition (Extended Data Fig. 8e) and retrieval of the associative memory was also similar between groups (Fig. 5e and Extended Data Fig. 8f).

These results further underline the sensitivity of enduring memory formation to changes in the excitation state of ACC engram ensembles during the early phase of memory consolidation.

## Discussion

We present findings showing that neuron-wide IE plasticity constitutes a compelling necessary and permissive mechanism for driving the time-dependent process of memory engram reorganization in the neocortex and controlling the informational richness of consolidated engrams. Neuronal allocation has been defined as the selection process that determines which specific neurons become parts of a given memory engram. Neurons exhibiting relatively higher excitability at the time of event encoding predominate in the selection process for engram allocation ^20^. In agreement with this concept, we revealed that learning-induced IE plasticity of a set of neurons in the ACC does occur shortly after associative encoding. This form of early neuronal tuning switches their state into a receptive (or “consolidating”) mode compatible with information processing and integration. In this mode, ACC engram neurons are more likely to be reactivated by the offline coordinated activity of hippocampal-cortical circuits during periods of quiet wakefulness or phases of sleep ^21^. This would promote their permanent integration into the cortical engram by driving persisting changes in synaptic and wiring plasticity ^22–24^, thereby strengthening the connections between nascent engram neurons undergoing consolidation-dependent maturation. Moreover, this increased excitability would prime the neurons to undergo further plasticity, for instance, by altering the associativity and cooperativity of synaptic inputs required to induce synaptic plasticity required for enduring storage.

When examining the learning-induced dynamics of IE plasticity, we discovered that the ‘tagged’ neurons in the ACC not only preferentially matured during the memory consolidation process (see Fig. 1E), leading to their strong recruitment upon remote memory retrieval but also that their IE underwent neuron-wide plastic modifications that were transient, occurring during the early, but not late, phase of memory storage. How then does this form of plasticity contribute to the remote expression of fear memory? This question has dogged discussions of how cortical IE plasticity could contribute to an enduring memory trace ^5,6,16^. As previously envisioned, early neuron-wide IE plasticity can set the stage for metaplasticity, for example, through gene-expression patterns mediated by ion-channel signaling ^25^, leading to downstream events that reinforce the communication within the nascent engram network. This is consistent with findings of early tagging events in the prefrontal cortex affecting the remote expression of non-aversive memories ^10^. In a landscape of increased excitability, synaptic inputs to specific neuronal compartments, such as dendritic branches, could create opportunities for localized excitability mechanisms ^26^. Such localized mechanisms intersect with ‘traditional’ mechanisms of plasticity, such as long-term potentiation (LTP) ^26–28^, and significantly increase the capacity for storing specific memories ^18,29^ due to the extended permutations possible with such mechanisms. As a result, changes in IE plasticity elicited by an associative learning episode could modulate the threshold for induction of an online wave of LTP in the synapses of allocated ACC neurons which in turn contributes to establishing the selectivity of their neuronal firing to an aversive context, as previously reported for the hippocampus ^30^. Another way that neuron-wide IE plasticity could contribute during the early consolidation period is to set the stage for synapse-specific tuning by eliciting changes in the intracellular localization and targeting of ion channel complexes. Such changes are known to occur during development ^31^ or pathophysiological states (Kv2.1 and epilepsy ^32^), and are often due to the association of pore-forming subunits of an ion channel with accessory subunits that confer properties of localization. The IE plasticity changes reported here are likely due to the orchestrated effect of a spectrum of different ion channel types (notably Na^+^, Ca^2+^, and K^+^ channels), and many of these are susceptible to modulatory processes such as phosphorylation or association with accessory subunits that affect their properties ^33^.

Because enduring memory storage requires repeated neuronal activation during periods of quiet restfulness or phases of sleep and develops slowly over several consecutive days via changes in the wiring and strength of synaptic connections ^34,35^, we reasoned that neuron-wide IE plasticity should last long enough to support permanent storage mechanisms. Conversely, this form of plasticity affecting synaptic penetrance would not be a suitable permanent memory storage mechanism since it would severely lower the storage capacity of neurons compared to one where information is stored in a synapse-specific manner. In line with these predictions, we found that IE plasticity returned to naive pre-learning levels during the early phase of systems consolidation, indicating that neuron-wide IE plasticity is time-limited and no longer required once the physical and chemical engram alterations underlying permanent memory storage have been established.

By artificially manipulating the excitation state in those ACC neurons initially tagged during the learning event (nascent engram neurons), we established the necessity of precocious, but not late, cortical IE plasticity for recruiting nascent cortical memory engram cells and for initiating their subsequent maturation and stabilization. Functionally, such a time-limited tuning of the neuron-wide IE may serve as a key gateway for triggering early offline LTP as well as the additional waves of synaptic plasticity events recently identified as prerequisites for systems consolidation ^30^.

Surprisingly, early iDREADD-mediated hyperpolarization, but not eDREADD-induced depolarization, of tagged ACC engram ensembles was beneficial for remote memory formation (enhanced consolidation) and for improving context discrimination (more richly detailed informational content). We pinpointed changes in the excitability of neurons in layer 5 of the ACC engram network, but other cortical layers or neuron types, including various interneuron types, are also likely targeted by our “tet-tagging” strategy. Thus, interventions targeting the excitation of the overall ACC engrams could have outcomes that might seem counterintuitive at first consideration but are consistent with the notion of this heterogeneity. The most parsimonious explanation for this unexpected finding is that hyperpolarization could restrict the competitive advantage of dominant ACC engram neurons, which is derived from their heightened responsiveness due to increased excitability. Tilting the balance toward non-engram cells would hinder the preferential activation of dominant fear engram cells by new oncoming experiences and impede their reallocation to other neuronal ensembles, thereby maintaining their initial context-specific memory content as consolidation proceeds. In this way, IE plasticity would act as an essential gating mechanism ensuring the embedding of encoding specificity within the boundaries of the cortical engram network initially allocated to the encoded memory. Fear generalization to irrelevant contexts would then be minimized by authorizing the allocation of other non-engram cells to novel context associations, whether similar or conflicting ^36^.

The operating features of cortical IE plasticity we uncovered have functional implications for consolidation-based processes. First, IE plasticity controls the informational content (or precision) of remote memories, which we addressed by challenging mice with different contexts upon remote memory retrieval. Promoting the reactivation of only those memories that are the most relevant for the context at hand has been conceptualized in the form of an index embedded precociously within the initially tagged engram network that would aid in coordinating memory retrieval ^37^. Since the ACC constitutes a central hub within the set of interconnected cortical regions supporting remote memory ^12^, it stands as a likely candidate for hosting such a coordinating cortically-based index, which could rely on IE plasticity as its main supportive mechanism. In this view, the IE plasticity gating mechanism offers an inherent cortical system for integrating the richly detailed content of memories being processed until their stabilization.

Second, IE plasticity enables cortically-maturing engrams to cope with oncoming interfering events. Experiencing a second proximate event can disrupt the consolidation of a first one (retrospective interference) possibly by reversing learning-induced LTP, resulting in memory forgetting ^38^. Concurring with this, we impaired remote memory retrieval when administering retroactively an interfering event during the early phase of memory consolidation. Pharmacogenetic hyperpolarization of ACC engram neurons rescued this deficit, likely by preventing their reallocation into the new interfering engram network.

Third, the restricted temporal dynamics of IE plasticity during systems consolidation points to a division of labor between complementary mechanisms supporting memory integration and enduring storage. The observation that tempering with IE plasticity during the late phase of systems consolidation did not impact remote memory retrieval indicates that additional intrinsic excitability-independent mechanisms, potentially restricted to more specific areas of the neuron such as synapses, take place to ensure the full maturation of cortical engram cells, including their well-organized synaptic architecture and connectivity within the engram cell network. It is reasonable to assume that it is only under this late storage configuration that enduring cortical memory engrams will express their full vividness upon retrieval.

Here, we unraveled the dynamics of IE plasticity and its operating features in configuring cortical engram neurons, highlighting its crucial role in determining the fate and informational content of remote, consolidated memories. Whether these features of cortical IE plasticity are specific to the consolidation of contextual fear memories or can be generalized to other aversive or appetitive memory forms remains to be further explored. Our study proves that modulating the IE state during its plastic phase renders nascent engram neurons resilient to interference, ensuring their successful maturation as cortical engram neurons. Investigating whether these mechanisms are equally advantageous in counteracting memory forgetting over time by preventing cortical engrams to undergo de-maturation (resulting in retrieval failure or memory erasure) would be highly relevant in physiology and physiopathology. We show that IE plasticity constitutes a necessary, permissive step of memory engram formation, suggesting that intrinsic excitability, synaptic plasticity, and synaptic reorganization mechanisms are intricately related during the formation of enduring cortical engrams. These complementary mechanisms likely cooperate to ensure the malleable nature and physical manifestation of cortical engrams, and future efforts should be devoted to understanding how novel information can be incorporated into non-naive neuronal networks already hosting pre-existing engrams or associative neocortical mental schemas ^39^.

## Materials and Methods

### Ethical considerations

All animal procedures, including anesthesia and surgery, were conducted under the European guidelines. The experiments were approved by the animal care and use committees of the Université de Bordeaux and the Institut National de la Santé et de la Recherche Médicale (INSERM), France. The ethics protocol numbers allocated to these experiments were 9000 and 12267.

### Animals

Double transgenic Tet-Tag mice (Fos-tTA and Fos-EGFP; tetO-lacZ and tTA*) (Jax# 008344) were used for all the experiments that required long-term labeling of putative engram neurons: electrophysiological, DREADD and interference experiments. Single transgenic mice (Fos-tTA or Fos-EGFP) were used for the rest of the experiments. Adult mice of both genders were used for all the behavioral and electrophysiological experiments. As we observed no sex differences in our measures, data from both genders were pooled. Mice aged 8-10 weeks were used for the electrophysiology experiments, while 8–15-week-old mice were used for all other experiments.

Mice were bred in the animal facility of the Neurocentre Magendie, group-housed, and kept at a 12-hour light-dark cycle (light on from 7 am to 7 pm) in temperature-humidity-controlled rooms and with *ad libitum* access to water and doxycycline chow (40 mg/kg). Doxycycline chow was replaced with standard chow (no doxycycline) 48h before fear conditioning to open the temporal window for tagging and provided back immediately following the learning experience.

### Viral constructs and stereotaxic surgeries

Two weeks before contextual fear conditioning, the ACC area of Tet-Tag mice was stereotactically injected with one of the following adeno-associated viruses (AAVs): AAV1/2-TRE-hM3Dq- mCherry, AAV1/2-TRE-hM4Di-mCherry or AAV1/2-TRE-YFP. Thirty minutes before the surgery, mice received a subcutaneous injection of the analgesic buprenorphine solution (0.1 mg/kg, AXIENCE SAS). Mice were then deeply anesthetized with 4% isoflurane, placed in a stereotaxic frame, then switched to 2% isoflurane for the rest of the surgery. Before incising the skin on the skull, mice received a subcutaneous injection of a local anesthetic, lidocaine (0.1 mg/ml, AXIENCE SAS; 0.05-0.1 ml), at the site of the incision. Surgery commenced once the mice displayed no reflexes after paw pinch. Bilateral stereotaxic infusions of the AAV were targeted to the ACC using the following coordinates (relative to bregma) and infusion angle: AP +0.9 mm, ML ± 0.75 mm, DV -1.65 mm, 20° from a vertical line. 500 nl of AAV virus was infused into each site at a rate of 100 nl/min. The needle was slowly removed 5 min after completion of the infusion. Mice were given a two-week post-surgery recovery period before starting experiments. The plasmid pAAV-TRE-YFP were kindly provided by Dr. Togenawa (MIT, USA), and the adeno-associated virus was produced in our Neurocentre Magendie Institute.

### Behavioral testing

#### Contextual fear conditioning (CFC)

Before behavioral training, mice were habituated in a waiting room and handled daily (10–20 min) for five consecutive days. Contextual fear conditioning (CFC) consisted of a 3-min free-exploring phase of the conditioning chamber by each mouse (context A, scented with 70% ethanol), followed by five 2 s-long foot shocks (intensity 0.7 mA) with a shock interval of 30 s. Context A was a grey plastic chamber (Table 4; 26 cm long, 25 cm in-depth, 18 cm in height, Imetronic, Marcheprime, France) with a metal grid floor to deliver electric shocks and a transparent Perspex door. After the shocks, the animals remained in the chamber for another 30 seconds. A control group (CTX) was exposed to context A, also scented with 70% ethanol, for the same amount of time (five minutes and 40 seconds in total) but without receiving any foot shocks. For memory retrieval, mice were re-exposed to the conditioning chamber (context A) for 3 min without receiving foot shocks or to context B (chamber with an additional white, prism-like box with a plastic floor that was shaped like a triangle and contained no metal grids) in the same room (see Table 4 for detailed comparison of contexts A and B). To evaluate the strength of fear memory during different temporal phases of memory formation, we assessed the freezing time of each mouse in a given context (A or B). Freezing behavior was considered as the absence of any movement, except for respiratory-related movement.

Freezing time during memory recall was manually counted by two independent experimenters who were blind to the experimental conditions. The percentage of freezing was calculated as the measured freezing time divided by the three minutes of the retrieval period. We also calculated the freezing time during the 30 seconds that followed the last shock of CFC (% freezing after learning) as the basal freezing level for every mouse. Delta freezing (Δ freezing) was calculated as follows: Δ freezing = (% freezing during retrieval) – (% freezing after learning).

#### Pharmacogenetic manipulation

To modulate the excitation state of neuronal ACC engram ensembles transduced with AAV1/2- TRE-hM3Dq-mCherry (excitatory DREADD, eDREADD) or AAV1/2-TRE-hM4Di-mCherry (inhibitory DREADD, iDREADD), mice received Clozapin-N-oxide (CNO) or vehicle (saline, Sal) via intraperitoneal (*i.p.*) injection twice a day. To modulate the excitability during the early stages of memory consolidation, the first CNO/Sal *i.p.* injection was administered 6 hours after the CFC and continued daily for the subsequent seven days. To alter the excitability of engram neuron ensembles during the late memory consolidation, mice received the first CNO/Sal *i.p.* injection on day 14 after the CFC and then for the following seven consecutive days (until day 21 following CFC). Injections were delivered between 9 and 10 a.m. and 3 and 4 p.m. on a given day. Mice were returned to their home cages immediately after each *i.p.* injection.

CNO (TOCRIS) was dissolved in DMSO (Sigma-Aldrich) at a stock concentration of 5 mg/ml and stored at -20℃. Aliquots of CNO stock solution were thawed before the injections and diluted to a final concentration of 0.5 mg/ml with a 0.9% NaCl solution. DMSO was diluted in the same manner and served as control (vehicle). CNO (or vehicle) was injected *i.p.* at 5 mg/kg body weight.

We verified the expression of the DREADD receptors within the ACC area of interest by imaging mCherry-positive neurons in brain slices using confocal microscopy (described below). Only mice with mCherry expression circumscribed to the appropriate ACC area were considered for histological, behavioral, and statistical analyses.

#### Memory interference protocol

On day 1 (early memory interference, early-INT) or day 15 (late memory interference, late-INT) following CFC, mice were either allowed to explore a novel environment for two hours (interference group) or stayed in their home cage (no-interference control group). This novel environment consisted of a cage twice the size of the home cage, covered with black paper, with only one transparent side to see the distal cues on the wall. Toys were placed inside the cage and rearranged every 20 minutes to stimulate exploratory behavior. In addition, mice had access to a running wheel for an hour. Water and doxycycline food was available for the duration of the whole experiment. The influence of interference on memory recall was assessed 30 days after CFC by placing mice in the same context where they were conditioned (context A) for 3 minutes.

To probe how pharmacogenetic hyperpolarization of ACC engram neurons alters the interference-induced impairment of memory formation, animals were injected with AAV1/2-TRE-hM4Di- mCherry into the ACC two weeks before CFC. Mice of the interference group were randomly divided into two subgroups, one receiving *i.p.* CNO injections and the other *i.p.* vehicle injections for seven days each during either the early- or remote phase of memory consolidation (see above). The no-interference control group received vehicle injections during the same 7-day period and stayed in their home cage. Memory retrieval was performed for all three groups 30 days after CFC in Context A.

### Electrophysiology

#### ACC slice preparation

Mice were sacrificed for electrophysiological recordings 1-3 days or 15-17 days following the behavioral session (CFC or CTX) without memory recall. Mice were deeply anesthetized with isoflurane, and intracardially perfused with ice-cold (4°C) cutting solution (for composition, see Table 5) once the mice displayed no reflexes after paw pinch. Following the perfusion, mice were decapitated, heads were immersed in the ice-cold cutting solution, and the brains were rapidly removed. Coronal slices of 300 µm were cut using a vibratome (Vibratome 3000 Plus, Sectioning Systems) and gently transferred to an incubating chamber filled with artificial cerebrospinal fluid (aCSF) resting solution (Table 5). Following 40 min of incubation at 37°C, slices were transferred to a second incubation chamber containing aCSF recording solution at room temperature (∼23℃) (Table 5). Brain slices were allowed to recover for at least one hour before electrophysiology recordings. All the solutions were saturated with carbogen (95% O2, 5% CO2), the pH adjusted to 7.32–7.40, and the osmolarity adjusted to 295–310 mOsm.

#### Whole-cell patch-clamp recordings

Following the behavioral protocol, mice were randomly assigned to the 1-3-day and 15-17-day groups for electrophysiological experiments. Layer 5 thick-tufted pyramidal neurons within acute ACC slices were selected for whole-cell recordings using an upright microscope (Zeiss Examiner) equipped with a 63X/1.0 NA water immersion objective (Zeiss) and an IR-Dodt contrast system. An Evolve 512 EMCCD Camera (Photometrics) combined with an LED system (560 nm; Colibri, Zeiss) and a Filter Set 62 HE (Zeiss) was used for visualizing YFP-expressing putative ACC engram neurons. For whole-cell recordings, borosilicate glass capillaries (Harvard Instruments, GC150F-7.5) were pulled using a PC-100 vertical puller (NARISHIGE Group). The pipettes were filled with internal recording solution (Table 5), and the open-tip resistance ranged from 5 to 7 MΩ. The osmolarity of the internal solution containing 0.3% biocytin (Reference: 90055, Biotium) was adjusted to 290 mOsm, and the pH was set to 7.30. The solution was stored at -20℃ and kept on ice before use. The recording chamber of the electrophysiology setup was perfused with oxygenated aCSF at a temperature of 32℃ during the whole experiment.

Recordings were performed in the bridge mode using a BVA-700C amplifier (Dagan, USA). Data was low-pass filtered at 3 kHz and sampled at 20 kHz using an ITC-18 (InstruTECH) Data Acquisition Interface and AxoGraph X software (version 1.7.6). Bridge resistance and capacitance were compensated and monitored during the recording period. Cells were discarded if the resting membrane potential was depolarized above -50 mV. All electrophysiological properties were measured in the bridge-mode.

To measure the intrinsic properties of ACC neurons, recordings were performed in the presence of the following blockers of glutamatergic and GABAergic synaptic transmission: AMPA receptor antagonist, NBQX (disodium salt, TOCRIS; concentration: 3μM in aCSF), NMDA receptor antagonist, DL-AP5 (TOCRIS; 50μM), and GABAA receptor antagonist, SR 95531 (hydrobromide, TOCRIS; 10μM).

#### Morphological reconstruction of recorded neurons

After electrophysiological recordings, the brain slices containing the biocytin-filled recorded neurons were fixed in 4% PFA for two hours and then transferred to 0.1 M PBS solution at 4℃. Brain slices were washed three times with 0.1 M PBS and then submerged for two hours at room temperature with a 0.7% Triton X-100 solution (EUROMEDEX). Slices were then incubated with a solution that contained Streptavidin-Alexa 555 (1:1000, Thermo Fisher, Reference: S21318) and 0.3% Triton X-100 for three hours. The brain slices were then washed with 0.1 M PBS three times and incubated with DAPI (1:10000, Sigma-Aldrich) for five minutes. Brain slices were mounted on microscope slides using Mowiol-Dabco and thin coverslips.

Confocal microscopy was used to image the recorded neurons (YFP+/Alexa 555+neurons). The distances of the recorded neurons to the pia and the beginning of layer 2 were also measured to define the layer localization of the neurons. The morphology of soma and dendrites was reconstructed with Neurolucida software, and only *post hoc* identified layer 5 tick-tufted pyramidal neurons were included for analysis.

#### Electrophysiological data analysis

Electrophysiological data were analyzed using AxoGraph X and customized Python analysis routines. The resting membrane potential (RMP) was measured 3 minutes after whole-cell access and switching to the bridge mode to allow for the cells to stabilize. The input resistance was measured as the average (five trials) steady-state voltage deflection to a -20 pA current step (800 ms duration). Rheobase and the number of action potentials (AP) as a function of current were determined by injecting a series of depolarizing current steps of increasing amplitude (between 20 and 300 pA, 20 pA increments, 800 ms duration). The rheobase was defined as the minimum amount of current necessary to evoke the first AP. The first AP within a train of 4-5 APs was analyzed for AP threshold (dV/dt exceeding 10mV/ms, in mV), AP amplitude (from RMP to peak amplitude, mV), AP half-width (duration at half-distance between AP threshold and amplitude, ms), and maximum rate of rise (dV/dt, mV/ms). The inter-spike interval (ISI, ms) was measured between the first two and the last two APs within a train of 4-5 APs, and the AP adaptation index was measured as follows: (first ISI) / (last ISI). The % of sag response was calculated using hyperpolarizing current injections as follows: (voltage difference between steady-state and the minimum of the hyperpolarization) / (voltage difference between the RMP and the minimum of the hyperpolarization) x 100 (in %). The membrane time constant (ms) was calculated from voltage responses to alternating depolarizing and hyperpolarizing 400 pA 1-ms-long current pulses. It was then calculated as the slow component of a double-exponential fit of the average voltage decay in both the depolarizing and hyperpolarizing directions. The medium afterhyperpolarization (mAHP) was measured as the minimum (relative to RMP) 50-100 ms following the last AP within a train of 5 APs (2 ms pulses, 2 nA) evoked at frequencies ranging from 20 to 120 Hz (at 20 Hz intervals), and 200 Hz. The slow AHP (sAHP) was measured as the minimum following the last AP in the train of 15 APs induced by 2 nA pulses at 20 Hz.

### Immunohistochemistry and imaging

#### Sacrifice and slice preparation

Ninety min after the behavior, mice were deeply anesthetized by 4% isoflurane and *i.p.* injected with a solution containing 400 mg/ml of pentobarbital (Exagon, Med’Vet) and 0.1 mg/ml of lidocaine (Lidor, AXIENCE SAS) in a 0.9% NaCl solution. The injection dose was 0.1 ml/10g of mouse body weight. Mice were then transcardially perfused with 0.1 M phosphate-buffered saline (PBS) followed by 4% cold paraformaldehyde (PFA). Mice of the home-cage group, which did not undergo any behavioral procedure, were perfused on the same day. Brains were dissected and postfixed overnight in 4% paraformaldehyde, then sliced into 50-μm coronal sections using a vibratome.

#### Immunohistochemistry staining

Brain slices were washed three times with 0.1 M PBS for 10 minutes each, and then blocked for one hour at room temperature with the following blocking solution: 5% BSA (EUROMEDEX) mixed with 0.3% Triton X-100 (EUROMEDEX). To enhance the GFP signal, slices were incubated with anti-GFP rabbit polyclonal antibody Alexa-488 (1:1000, Thermo Fisher, Reference: A21311) overnight at 4℃ in a solution containing 1% BSA and 0.3% Triton X-100. The following day, slices were washed three times with 0.1 M PBS for 10 minutes each and incubated with DAPI (1:10000, Sigma-Aldrich) for 5 minutes. After being washed once with 0.1 M PBS, the brain slices were mounted using Mowiol-Dabco and thin coverslips on microscope slides.

#### Confocal image acquisition and analysis

Fluorescence images were acquired using a Zeiss LSM700 confocal microscope equipped with a Leica 20X/1.3 NA oil immersion lens provided by the Bordeaux Imaging Centre (BIC). Two to three coronal slices per mouse were imaged at multiple depths along the z-axis (z-stack of 32 µm with 4 µm interval steps). Imaris software (version 9.6.2) enabled the reconstruction of a 3- dimensional model from confocal microscopy images. Cell counting analysis was performed using Imaris and ImageJ-Fiji software (provided by BIC). The area of the ACC was selected and measured with ImageJ-Fiji. The number of mCherry-positive, GFP-positive, co-localized (mCherry+/GFP+), and DAPI-positive neurons were counted and expressed as density per mm2 ((#cells/area size (um2)) x 1000000). Image acquisition and analysis were performed blind to the experimental conditions.

### Statistical Analysis

Statistical tests were performed using Prism 10.0 (GraphPad). The normality distribution of the data was examined using the D’Agostino-Pearson omnibus normality test. If the data were normally distributed, an unpaired Student’s *t-test*, a one-way ANOVA, or a two-way ANOVA test was performed. Otherwise, nonparametric tests were applied. When performing the ANOVA test, the standard deviation of the data set was examined. To identify potential outliers the ROUT (Q=0.1%) method was used. The post-hoc multiple comparisons were performed using the Tukey or Šídák’s test. Data were displayed as mean ± SEM. The significance threshold was set at α=0.05 (NS, *\* P* < 0.05*, ** P* < 0.01*, ***P* < 0.001*, **** P* < 0.0001).

## Resource availability

Further information and requests for resources should be directed to and will be fulfilled by the lead contact, Andreas Frick

## Materials availability

All unique/stable reagents generated in this study are available from the corresponding authors upon request.

## Data and code availability

All data and code are available in the manuscript, the supplementary material or at the request to the corresponding authors.

## Acknowledgments

We thank past and current members of the Frick and Bontempi laboratories for fruitful discussions and comments that contributed to the progress of this project. We thank Elodie Cougouilles for assistance with behavioral experiments and Francesca Bettoni for assistance with neuronal reconstructions. We thank Dr. Ourania Semelidou and Dr. Marie Oulé for feedback on the manuscript. We thank Dr. Aline Desmedt for providing access to her behavioral equipment. We thank the genotyping and animal housing facilities of INSERM U1215 Neurocentre Magendie for ensuring the welfare of the laboratory animals. We would like to acknowledge the Bordeaux Imaging Center, a service unit of CNRS-INSERM and Bordeaux University and member of the national infrastructure France BioImaging supported by the French National Research Agency (ANR-10-INBS-04), and Dr. Monica Fernandez Monreal and Sebastien Marais for their technical assistance with microscopy and image analysis. We thank Dr. Susumu Togenawa (MIT) for kindly providing the plasmids for activity-dependent neuronal labeling.

## Funding

Agence Nationale de la Recherche grant (AF, BB)

The LabEx BRAIN funding (BB, AF)

INSERM endowment (AF)

China Scholarship Council (LZ)

University of the Basque Country - PIFBUR/20 (MGC)

## Author contributions

Conceptualization: SH, MG, BB, AF

Experiments: SH, LZ, MG, RDS, MGC, KLC, ON

Analysis: SH, LZ, YLF

Funding acquisition: LZ, MGC, BB, AF

Project administration: BB, AF

Writing – original draft: SH, LZ, MG, BB, AF

Writing – review & editing: SH, LZ, MG, YLF, ON, BB, AF

## Diversity, equity, ethics, and inclusion

We support inclusive, diverse, and equitable conduct of research. All experimental procedures were conducted following the official European Guidelines for the Care and Use of Laboratory Animals (86/609/EEC), approved by the Bordeaux ethics committee under the aegis of the French Ministry of Research, and followed the ARRIVE guidelines.

## Extended Data figures include

Figures 1-8

Tables 1-5

### Extended Data figure titles and legends

**Extended Data Fig. 1. Contextual fear conditioning (CFC) training data related to** Fig. 1c and f**. a,** Freezing behavior, expressed as % freezing, following the last shock of CFC was similar between groups that would subsequently undergo retrieval at 1 and 30 days, indicating comparable levels of acquisition. Data points are individual mice with mean ± SEM. Statistical significance was calculated using the unpaired t-test.

**Extended Data Fig. 2. Contextual fear conditioning triggers IE plasticity in ACC nascent engram ensembles. a,** Experimental design: whole-cell patch-clamp recordings were performed from from neurons expressing YFP (YFP^+^) 1-3 days post-CFC or post-CTX. **B,** Trend for reduced AP adaptation in CFC-YFP^+^ neurons (*P* = 0.108). **C,** Increase in Tau in CFC-YFP^+^ neurons. Data points are individual cells with median, lower and upper quartiles, and minimum and maximum data values. Statistical significance was calculated using the unpaired t-test, \*\*\**P* < 0.001.

**Extended Data Fig. 3. CFC-induced IE plasticity of ACC nascent engram ensembles waned during the remote memory phase. a,** Experimental design: whole-cell patch-clamp recordings were performed from YFP^+^ neurons 15-17 days post-CFC or post-CTX. **b-l**, Analysis for n=12 YFP^+^ cells (CFC group) and n=12 YFP^+^ cells (CTX group). No significant difference was found for the following electrophysiological parameters: (**b**) input resistance, (**c**) number of APs as a function of injected current (0-300 pA), (**d**) RMP, (**e**) Rheobase, (**f**) AP amplitude, (**g**) AP halfwidth, (**h**) AP threshold, (**i**) AP max dv/dt, (**j**) Tau depolarizing, (**k**) AP adaptation and (**l**) mAHP from a 5-AP-train at different frequencies. Data points are individual cells with median, lower and upper quartiles, and minimum and maximum data values. Statistical significance was calculated using repeated measures of two-way ANOVA followed by Šídák’s multiple comparisons tests (b, l), or the unpaired t-test (c-k).

**Extended Data Fig. 4. Transient nature of IE plasticity of the ACC engram neurons. a,** Experimental design: whole-cell patch-clamp recordings were performed from tagged (YFP^+^) L5 pyramidal neurons either during early-phase (1-3 days post-CFC) or remote-phase (15-17 days post-CFC) memory consolidation. Data are normalized to values from control cells (CTX-YFP^+^) at the respective memory phases. **b-f**, Analysis for n=9 cells (early-phase memory) and n=12 cells (remote-phase memory). Significant differences were found for the AP adaptation (**b**) and Tau depolarizing (**c**). No significant difference was observed for input resistance (**d**), AP max dv/dt (**e**), and a trend for an increased AP threshold (*P* = 0.084) during the early phase (**f**). Data points are individual cells with median, lower and upper quartiles, and minimum and maximum data values. Statistical significance was calculated using the unpaired t-test, \**P* < 0.05, \*\*\**P* < 0.001.

**Extended Data Fig. 5. Impact of context exposures on freezing behavior in the other context. a (**left), Experimental design. (right) When mice were tested in Context A in the morning session and Context B in the afternoon session, they were able to discriminate between the two contexts across all delays (1, 21, 30, and 40 days post-CFC). **b (**left), Experimental design. (right) When mice were tested in Context B in the morning session and Context A in the afternoon session, they were not able to discriminate between the two contexts across all delays (1, 21, 30, and 40 days post-CFC). **c (**left), Experimental design. Mice were tested only in Context A or Context B to avoid the impact of previous exposure. (right) Mice were able to discriminate between contexts across all delays, with significantly increased levels of freezing between day 1 and day 40 for Context B. Data points are individual mice with mean ± SEM. Statistical significance was calculated using two-way ANOVA followed by Tukey/Šídák’s multiple comparisons test (a-c), \**P* < 0.05, \*\**P* < 0.01, **** *P* < 0.0001.

**Extended Data Fig. 6. CFC training data, ACC engram labeling during encoding, and retrieval activation following pharmacogenetic manipulation during early-phase memory consolidation. a,** Experimental design. **b,** Training data related to Fig. 4c. Results are presented for the three groups (iDREADD/CNO, eDREADD/CNO, i/eDREADD/Sal) as the % of time spent freezing during one minute following the last shock of CFC, showing no difference between groups. **c**, Density of mCherry^+^ cells does not differ between groups, suggesting similar number of engram neurons that expressed DREADD. **d,** Density of GFP^+^ cells indicates an increase in number of cells that were active during the retrieval in Context B, but no effect of treatment. Data points are individual mice with mean ± SEM. Statistical significance was calculated using two-way ANOVA followed by Tukey/Šídák’s multiple comparisons test (b-d).

**Extended Data Fig. 7. CFC training data, ACC engram labeling during encoding, and retrieval activation following pharmacogenetic manipulation during remote-phase memory. a,** Experimental design. **b**, Training data related to Fig. 4G. Results are presented for the three groups (iDREADD/CNO, eDREADD/CNO, i/eDREADD/Sal) as the % of time spent freezing during one minute following the last shock of CFC, showing no difference between groups. **c,** Density of mCherry^+^ cells does not differ between groups, suggesting a similar number of engram neurons that expressed DREADD. **d**, Density of GFP^+^ cells labeling neurons active during retrieval shows no difference among the groups. Data points are individual mice with mean ± SEM. Statistical significance was calculated using two-way ANOVA followed by Tukey/Šídák’s multiple comparisons test (b-d).

**Extended Data Fig. 8.** CFC training data and neuronal activation during encoding and retrieval in interference experiments. a, Experimental design. b, Training data related to Fig. 5a and b. Results are presented for the three groups (no-INT, Early INT, Late INT) as the % of time spent freezing during one minute following the last shock of CFC, showing no difference between groups. c, Experimental design. d, Training data related to Fig. 5c to e. Results are presented for the three groups (no-INT Sal, INT Sal, INT CNO) as the % of time spent freezing during one minute following the last shock of CFC, showing no difference between groups. e, Density of mCherry^+^ cells does not differ between groups, suggesting a similar number of engram neurons that expressed DREADD. f, Density of GFP^+^ cells labeling neurons active during retrieval shows no difference among the groups. Data points are individual mice with mean ± SEM. Statistical significance was calculated using two-way ANOVA followed by Tukey/Šídák’s multiple comparisons test (b, d-f).

