## Supplementary data for "Early intrinsic plasticity of neocortical engram neurons defines memory formation and precision"

**This file includes:**

Extended Data Fig. 1 to 8
Tables 1 to 5

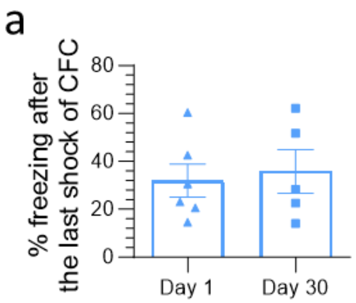


**Extended Data Fig. 1. Contextual fear conditioning (CFC) training data related to Fig. 1c and f. a,** Freezing behavior, expressed as % freezing, following the last shock of CFC was similar between groups that subsequently underwent retrieval at 1 and 30 days, indicating comparable levels of acquisition ($P$ = 0.74, NS). Data points are individual mice with mean ± SEM. Statistical significance was assessed using the unpaired t-test.

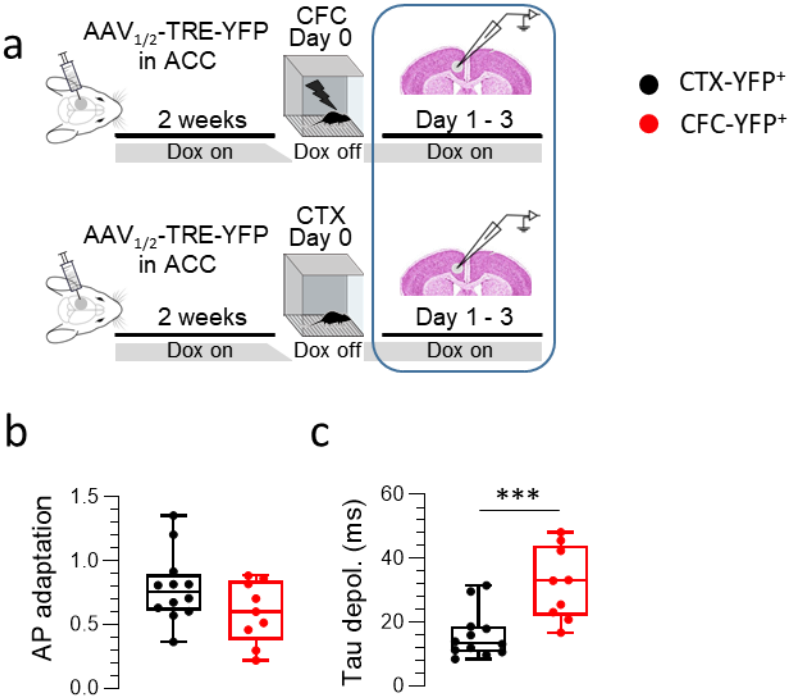


**Extended Data Fig. 2. Contextual fear conditioning triggers IE plasticity in ACC nascent engram ensembles.** Related to Fig. 2. **a,** Experimental design: whole-cell patch-clamp recordings were performed from neurons expressing YFP (YFP$^{+}$) 1-3 days post-CFC or post-CTX. **b,** Trend for reduced AP adaptation in CFC-YFP$^{+}$ neurons ($P = 0.108$, NS). **c**, Increase in Tau in CFC-YFP$^{+}$ neurons. Data points are individual cells with median, lower and upper quartiles, and minimum and maximum data values. Statistical significance was assessed using the unpaired t-test, $***P < 0.001$.

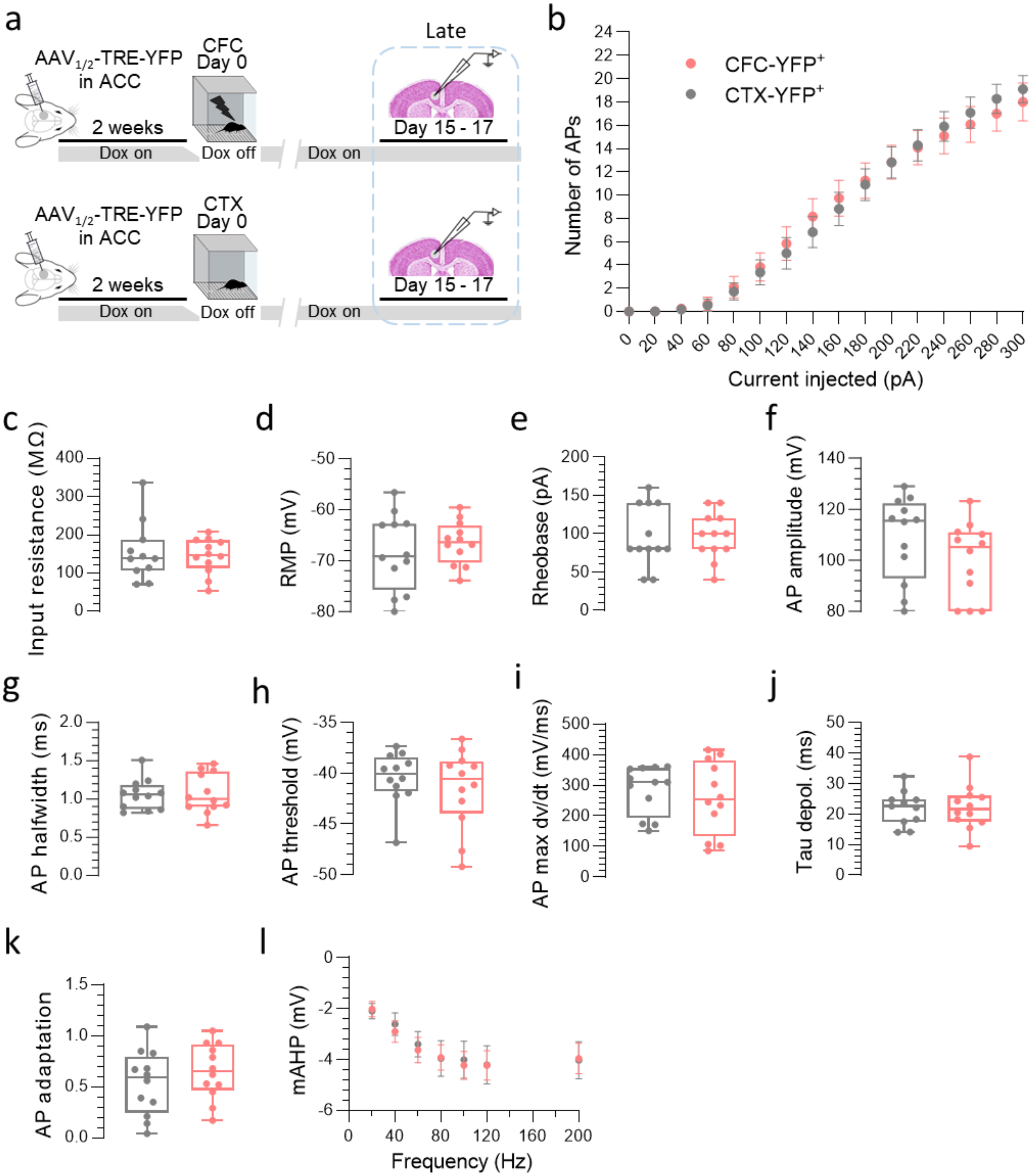


**Extended Data Fig. 3. CFC-induced IE plasticity of ACC nascent engram ensembles waned during the late phase of memory consolidation. a,** Experimental design: whole-cell patch-clamp recordings were performed from YFP$^+$ neurons 15-17 days post-CFC (in red) or post-CTX (in grey). **b-l**, Analysis for n = 12 YFP$^+$ cells (CFC group) and n = 12 YFP$^+$ cells (CTX group). No significant difference assessed by unpaired t-tests (c-k) or two-way ANOVA with repeated measures followed by Šídák's multiple comparisons tests (b and l) was found for the following electrophysiological parameters ($p > 0.05$ for all comparisons, NS): (**b**) Input resistance, (**c**) Number of APs as a function of injected current (0-300 pA), (**d**) RMP, (**e**) Rheobase, (**f**) AP amplitude, (**g**) AP halfwidth, (**h**) AP threshold, (**i**) AP max dv/dt, (**j**) Tau depolarizing, (**k**) AP adaptation and (**l**) mAHP from a 5-AP-train at different frequencies. Data points are individual cells with median, lower and upper quartiles, and minimum and maximum data values.

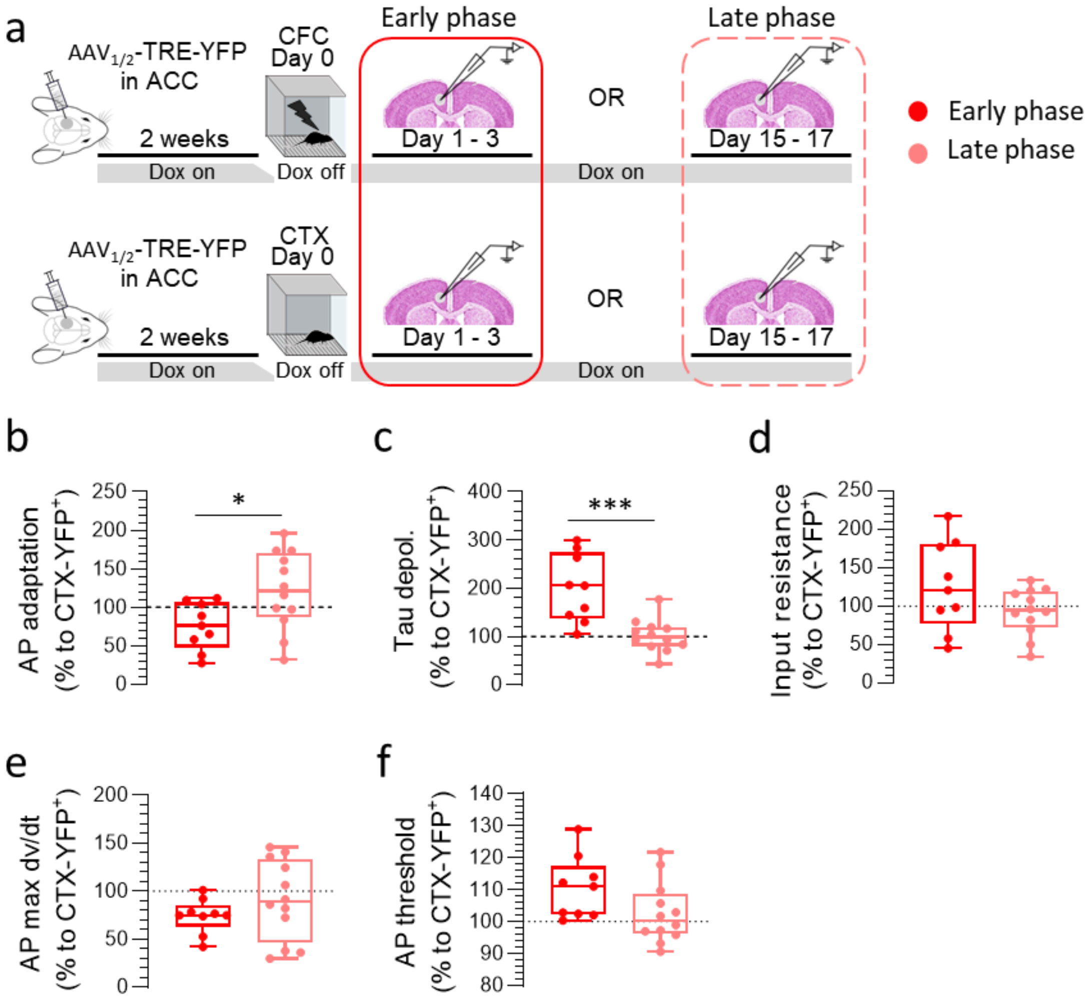


**Extended Data Fig. 4. Transient nature of IE plasticity of the ACC engram neurons.** Related to Fig. 3. **a,** Experimental design: whole-cell patch-clamp recordings were performed from tagged ($YFP^+$) L5 pyramidal neurons either during early-phase (1-3 days post-CFC) or remote-phase (15-17 days post-CFC) memory consolidation. Data are normalized to values from control cells (CTX-$YFP^+$) at the respective memory phases. **b-f,** Analysis for n = 9 cells (early phase of memory consolidation) and n = 12 cells (remote phase of memory consolidation). Significant differences were found for the AP adaptation (**b**) and Tau depolarizing (**c**). No significant difference was observed for input resistance (**d**), AP max dv/dt (**e**), and **only** a trend for an increase in the AP threshold ($P = 0.084$) during the early phase (**f**). Data points are individual cells with median, lower and upper quartiles, and minimum and maximum data values. Statistical significance was calculated using the unpaired t-test, *$P < 0.05$, ***$P < 0.001$.

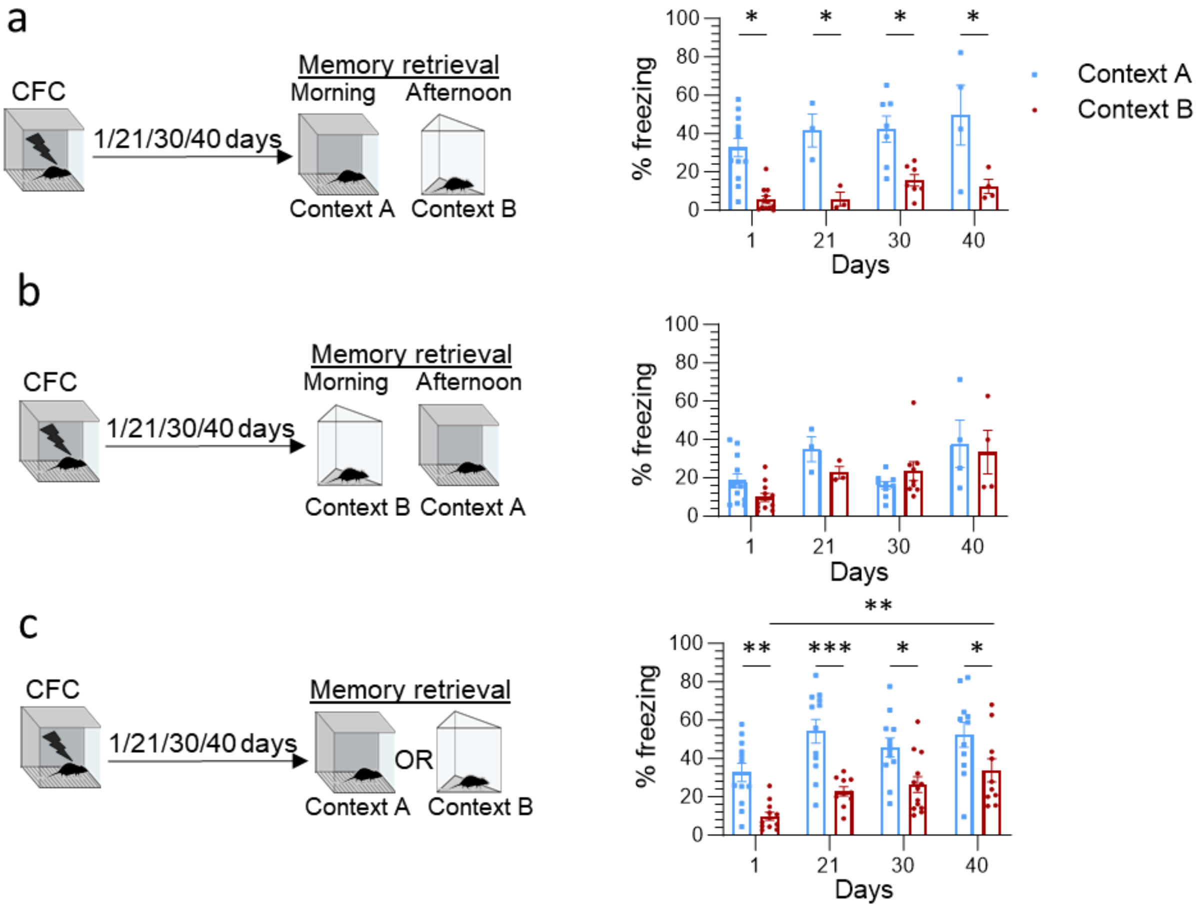


**Extended Data Fig. 5. Impact of context exposures on freezing behavior in the other context.** **a** (left), Experimental design. (right), When mice were tested in Context A in the morning session and Context B in the afternoon session, they were able to discriminate between the two contexts across all delays (1, 21, 30, and 40 days post-CFC). **b** (left), Experimental design. (right), When mice were tested in Context B in the morning session and Context A in the afternoon session, they were not able to discriminate between the two contexts across all delays (1, 21, 30, and 40 days post-CFC). **c** (left), Experimental design. Mice were tested only in Context A or Context B to avoid the impact of previous exposure. (right), Mice were able to discriminate between contexts across all delays, with significantly increased levels of freezing between day 1 and day 40 for Context B. Data points are individual mice with mean ± SEM. Statistical significance was calculated using two-way ANOVA followed by Tukey/Šídák's multiple comparisons test (a-c), *$P < 0.05$, **$P < 0.01$, **** $P < 0.0001$.

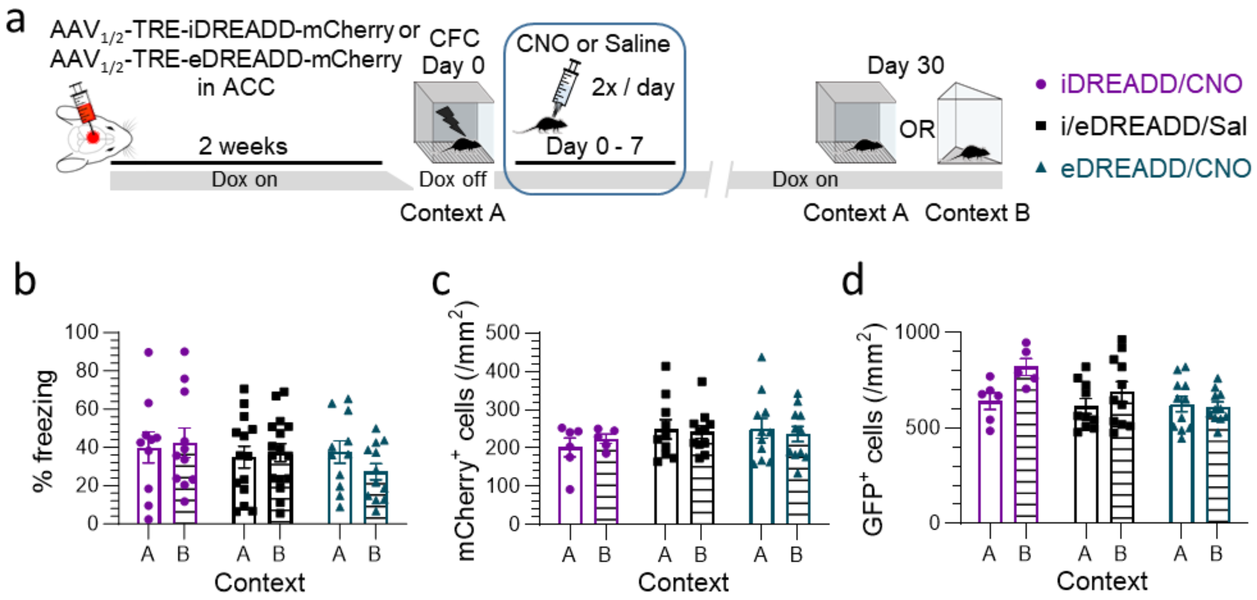


**Extended Data Fig. 6. CFC training data, ACC engram labeling during encoding, and retrieval activation following chemogenetic manipulation during the early phase of memory consolidation. a,** Experimental design. **b,** Training data related to Fig. 4c. Results are presented for the three groups (iDREADD/CNO, eDREADD/CNO, i/eDREADD/Sal) as the % of time spent freezing during one minute following the last shock of CFC, showing no difference between groups. **c**, Density of mCherry$^{+}$ cells does not differ between groups, suggesting that a similar number of engram neurons expressed DREADD. **d,** Density of GFP$^{+}$ cells indicates an increase in the number of cells that were active during the retrieval in Context B, but no effect of treatment. Data points are individual mice with mean ± SEM. Statistical significance was calculated using two-way ANOVA followed by Tukey/Šídák's multiple comparisons test (b-d).

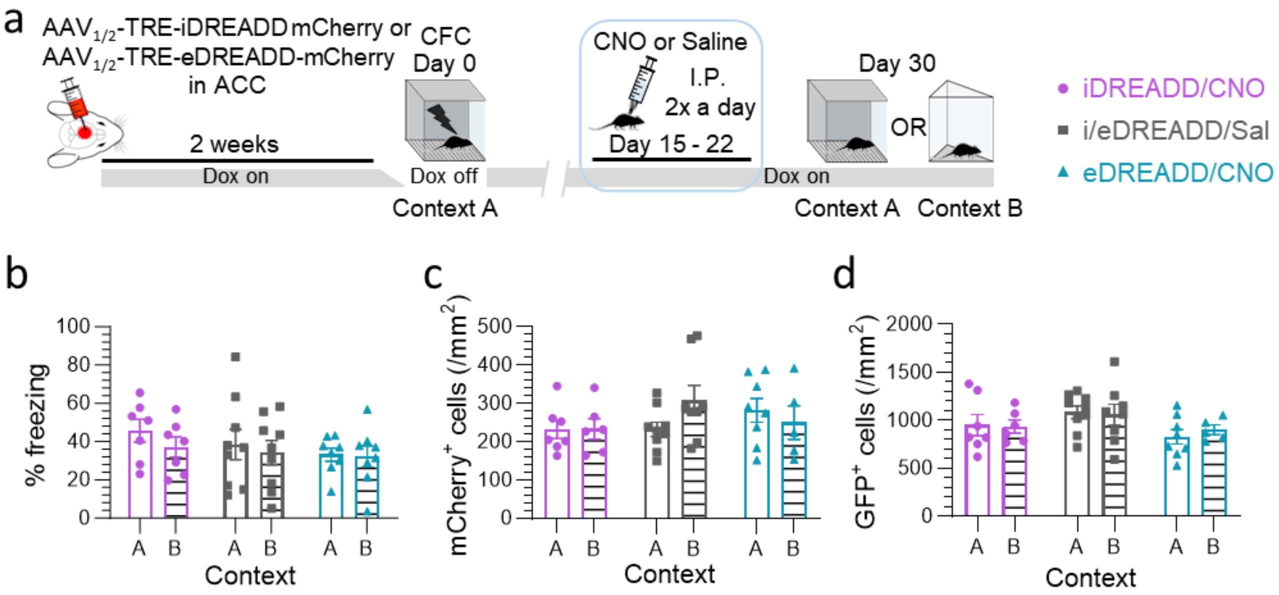


**Extended Data Fig. 7. CFC training data, ACC engram labeling during encoding, and retrieval activation following chemogenetic manipulation during the late phase of memory consolidation. a,** Experimental design. **b**, Training data related to Fig. 4g. Results are presented for the three groups (iDREADD/CNO, eDREADD/CNO, i/eDREADD/Sal) as the % of time spent freezing during one minute following the last shock of CFC, showing no difference between groups. **c,** Density of $mCherry^+$ cells does not differ between groups, suggesting that a similar number of engram neurons expressed DREADD. **d**, Density of $GFP^+$ cells is similar across groups, indicating that labeling of neurons active during retrieval was not a confounding factor. Data points are individual mice with mean ± SEM. Statistical significance was calculated using two-way ANOVA followed by Tukey/Šídák's multiple comparisons test (b-d).

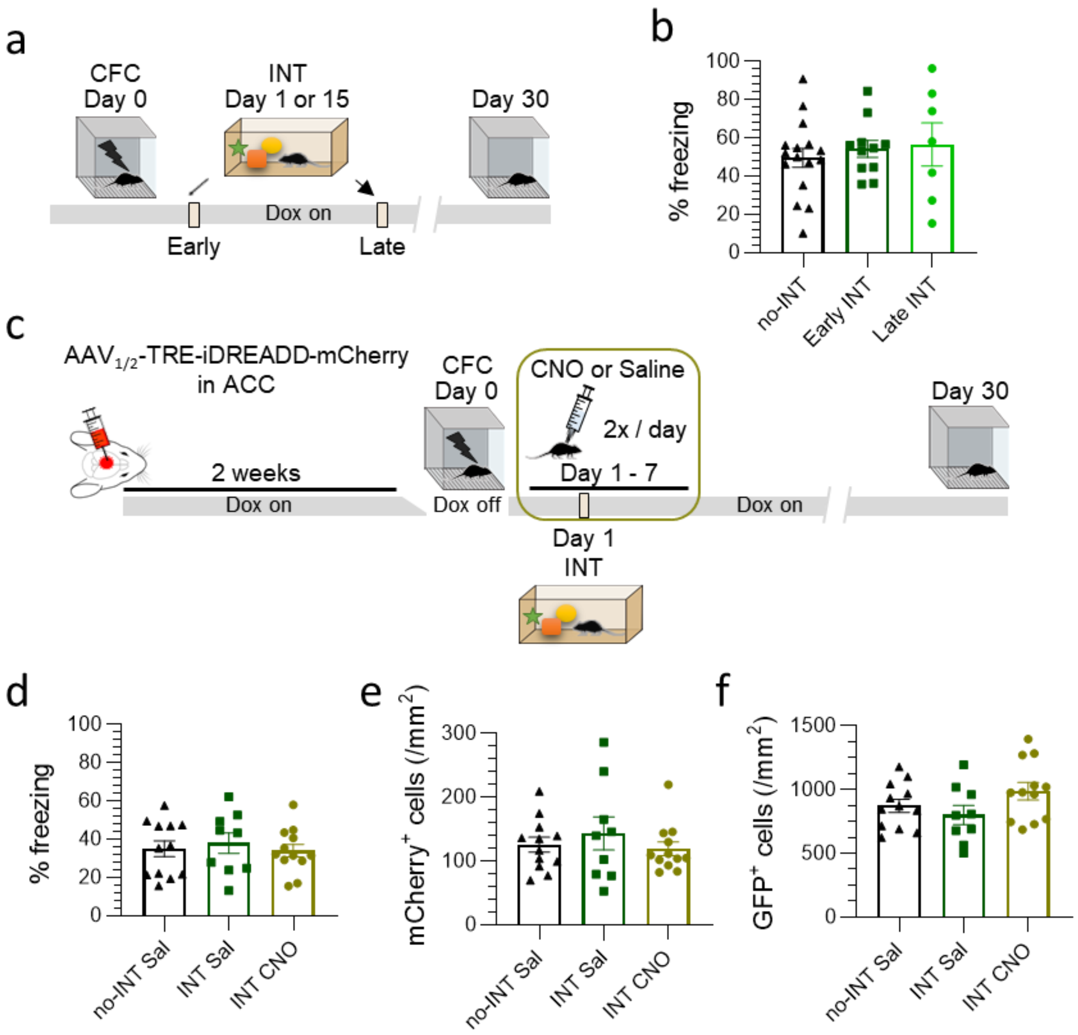


**Extended Data Fig. 8. CFC training data and neuronal activation during encoding and retrieval in experiments assessing the effects of interference. a,** Experimental design. **b,** Training data related to Fig. 5a and b. Results are presented for the three groups (no-INT, Early INT, Late INT) as the % of time spent freezing during one minute following the last shock of CFC, showing no difference between groups. **c,** Experimental design. **d,** Training data related to Fig. 5c to e. Results are presented for the three groups (no-INT Sal, INT Sal, INT CNO) as the % of time spent freezing during one minute following the last shock of CFC, showing no difference between groups. **e,** Density of mCherry$^{+}$ cells does not differ between groups, suggesting that a similar number of engram neurons expressed DREADD. **f,** Density of GFP$^{+}$ cells is similar across groups, indicating that labeling of neurons active during retrieval was not a confounding factor. Data points are individual mice with mean ± SEM. Statistical significance was calculated using two-way ANOVA followed by Tukey/Šídák's multiple comparisons test (b, d-f).

**Table 1. A complete list of intrinsic excitability measures and statistical analysis from whole-cell patch-clamp recordings performed from YFP$^{+}$ neurons 1-3 days post-CFC or post-CTX.**

| | Normality | Outlier | Test | | | P |
|---|---|---|---|---|---|---|
| **RMP** | Yes | No | Unpaired t test | | t19 = 2.32 | ***0.0318*** |
| **Tau dep** | Yes | No | Unpaired t test | | t19 = 3.90 | ***0.001*** |
| **Rheobase** | Yes | No | Unpaired t test | | t19 = 3.21 | ***0.0046*** |
| **Nb of APs** | Yes | No | 2way ANOVA | Interaction | F (15, 285) = 2.29 | ***0.0044*** |
| | | | | Current injected | F (15, 285) = 93.00 | ***<0.0001*** |
| | | | | Learning | F (1, 19) = 2.02 | 0.1716 |
| **mAHP after 5 AP-train** | Yes | No | Mixed-effects model (REML) | Interaction | F (6, 113) = 8.71 | ***<0.0001*** |
| | | | | Hz | F (6, 113) = 38.58 | ***<0.0001*** |
| | | | | Learning | F (1, 19) = 0.51 | 0.4822 |
| **AP threshold** | Yes | No | Unpaired t test | | t19 = 2.73 | ***0.0134*** |
| **AP halfwidth** | No | Yes (1) | Mann-Whitney | | U = 8 | ***0.0004*** |
| **AP max dv/dt** | Yes | No | Unpaired t test | | t19 = 2.15 | ***0.0446*** |
| **AP amplitude** | Yes | No | Unpaired t test | | t19 = 2.17 | ***0.0431*** |
| **AP adaptation** | Yes | No | Unpaired t test | | t19 = 1.68 | *0.1085* |
| **Input resistance** | Yes | No | Unpaired t test | | t19 = 1.18 | 0.2515 |
| **Sag ratio** | Yes | No | Unpaired t test | | t19 = 0.41 | 0.6876 |
| **sAHP after 15 AP-train** | No | No | Mann-Whitney | | U = 40 | 0.3451 |

Significant P values are shown in bold.

**Table 2. A complete list of intrinsic excitability measures and statistical analysis from whole-cell patch-clamp recordings performed from YFP+ neurons 15-17 days post-CFC or post-CTX.**

| | Normality | Outlier | Test | | | P |
|---|---|---|---|---|---|---|
| **RMP** | Yes | No | Unpaired t test | | t22 = 0.76 | 0.4541 |
| **Tau dep** | Yes | No | Unpaired t test | | t21 = 0.11 | 0.9128 |
| **Rheobase** | Yes | No | Unpaired t test | | t21 = 0.00 | >0.9999 |
| **Nb of APs** | Yes | No | 2way ANOVA | Interaction | F (15, 315) = 0.55 | 0.9106 |
| | | | | Current injected | F (15, 315) = 186.0 | ***<0.0001*** |
| | | | | Learning | F (1, 21) = 0.00 | 0.9965 |
| **mAHP after 5 AP-train** | Yes | No | Mixed-effects model (REML) | Interaction | | 0.7913 |
| | | | | Hz | | ***<0.0001*** |
| | | | | Learning | | 0.7445 |
| **AP threshold** | Yes | No | Unpaired t test | | t22 = 0.42 | 0.4187 |
| **AP halfwidth** | Yes | No | Unpaired t test | | t22 = 0.13 | 0.8982 |
| **AP max dv/dt** | Yes | No | Unpaired t test | | t22 = 0.65 | 0.5231 |
| **AP amplitude** | Yes | No | Unpaired t test | | t22 = 1.51 | 0.1446 |
| **AP adaptation** | Yes | No | Unpaired t test | | t22 = 0.96 | 0.3469 |
| **Input resistance** | Yes | No | Unpaired t test | | t21 = 0.3974 | 0.6951 |
| **Sag ratio** | Yes | No | Unpaired t test | | t22 = 0.17 | 0.8647 |
| **sAHP after 15 AP-train** | Yes | No | Unpaired t test | | t20 = 1.82 | *0.0842* |

Significant P values are shown in bold.

**Table 3. Statistical comparison of intrinsic excitability measures from YFP⁺ neurons 1-3 days and 15-17 days post-CFC normalized to those of CTX during early and late phases of memory consolidation, respectively.**

| | Normality | Outlier | Test | | | P |
|---|---|---|---|---|---|---|
| **RMP** | Yes | No | Unpaired t test | | t19 = 2.75 | ***0.0127*** |
| **Tau dep** | Yes | No | Unpaired t test | | t19 = 4.24 | ***0.0004*** |
| **Rheobase** | Yes | No | Unpaired t test | | t19 = 3.40 | ***0.003*** |
| **Δ Nb of APs** | Yes | No | 2way ANOVA | Interaction | F (15, 285) = 2.60 | ***0.0011*** |
| | | | | Current injected | F (15, 285) = 5.29 | ***<0.0001*** |
| | | | | Learning | F (1, 19) = 2.15 | 0.1591 |
| **Δ mAHP after 5 AP-train** | Yes | No | Mixed-effects model (REML) | Interaction | F (6, 113) = 5.94 | ***<0.0001*** |
| | | | | Hz | F (6, 113) = 8.94 | ***<0.0001*** |
| | | | | Learning | F (1, 19) = 0.41 | 0.5274 |
| **AP threshold** | Yes | No | Unpaired t test | | t19 = 1.82 | *0.084* |
| **AP halfwidth** | Yes | No | Unpaired t test | | t19 = 6.70 | ***<0.0001*** |
| **AP max dv/dt** | Yes | No | Unpaired t test | | t19 = 1.13 | 0.2734 |
| **AP amplitude** | Yes | No | Unpaired t test | | t19 = 2.89 | ***0.0094*** |
| **AP adaptation** | Yes | No | Unpaired t test | | t19 = 2.41 | ***0.0263*** |
| **Input resistance** | Yes | No | Unpaired t test | | t19 = 1.68 | *0.1088* |
| **Sag ratio** | Yes | No | Unpaired t test | | t19 = 0.62 | 0.5415 |
| **sAHP after 15 AP-train** | No | No | Mann-Whitney | | U = 38 | 0.2773 |

Significant P values are shown in bold.

**Table 4. Features of Context A and Context B.**

| | Context A | Context B |
|---|---|---|
| **Photo** | 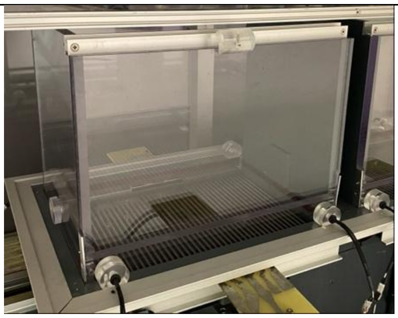 | 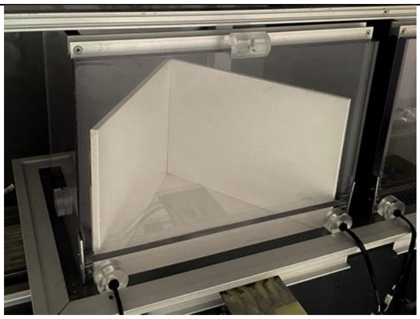 |
| **Size** | L:26 cm, D: 24cm, H:18 cm | L:24 cm, D:17 cm, H:15 cm |
| **Floor** | Metal grid, rectangular | Plastic pad, triangle |
| **Wall** | Grey plastic wall with one transparent door | White plastic wall with one transparent door |
| **Odor** | 70% ethanol | No ethanol |
| **Light** | Ambient light | Ambient light |

**Table 5. Composition of external and pipette solutions.**

| Ingredients (mM) | Cutting solution | aCSF resting solution | aCSF recording solution | Internal solution |
|---|---|---|---|---|
| **NaCl** | - | 124 | 124 | - |
| **NMDG** | 93 | - | - | - |
| **KCl** | 2.5 | 2.5 | 2.5 | 4 |
| **$NaH_2PO_4 \cdot H_20$** | 1.25 | 1.25 | 1.25 | - |
| **$NaHCO_3$** | 30 | 24 | 24 | - |
| **HEPES** | 20 | 5 | 5 | 10 |
| **D-Glucose** | 35 | 12.5 | 12.5 | - |
| **Thiourea** | 2 | - | - | - |
| **Na-ascorbate** | 5 | 1 | - | - |
| **Na-pyruvate** | 3 | 4 | - | - |
| **$CaCl_2 \cdot 2H_2O$** | 0.5 | 2 | 2 | - |
| **$MgSO_4 \cdot 7H_2O$** | 10 | 2 | 2 | - |
| **K-gluconate** | - | - | - | 135 |
| **Phospocreatine-Na** | - | - | - | 10 |
| **Na-GTP** | - | - | - | 0.3 |
| **Mg-ATP** | - | - | - | 4 |
